# Dopamine role in learning and action inference

**DOI:** 10.1101/837641

**Authors:** Rafal Bogacz

## Abstract

This paper describes a framework for modelling dopamine function in the mammalian brain. In this framework, dopaminergic neurons projecting to different parts of the striatum encode errors in predictions made by the corresponding systems within the basal ganglia. These prediction errors are equal to differences between rewards and expectations in the goal-directed system, and to differences between the chosen and habitual actions in the habit system. The prediction errors enable learning about rewards resulting from actions and habit formation. During action planning, the expectation of reward in the goal-directed system arises from formulating a plan to obtain that reward. Thus dopaminergic neurons in this system provide feedback on whether the current motor plan is sufficient to obtain the available reward, and they facilitate action planning until a suitable plan is found. Presented models account for dopaminergic responses during movements, effects of dopamine depletion on behaviour, and make several experimental predictions.

## Introduction

Neurons releasing dopamine send widespread projections to many brain regions, including basal ganglia and cortex (Björklund & Dunnett, 2007), and substantially modulate information processing in the target areas. Dopaminergic neurons in the ventral tegmental area respond to unexpected rewards (Schultz, Dayan, & Montague, 1997), and hence it has been proposed that they encode reward prediction error, defined as the difference between obtained and expected reward (Houk, Adams, & Barto, 1995; Montague, Dayan, & Sejnowski, 1996). According to the classical reinforcement learning theory, this prediction error triggers update of the estimates of expected rewards encoded in striatum. Indeed, it has been observed that dopaminergic activity modulates synaptic plasticity in the striatum in a way predicted by the theory (Reynolds, Hyland, & Wickens, 2001; Shen, Flajolet, Greengard, & Surmeier, 2008). This classical reinforcement learning theory of dopamine has been one of the greatest successes of computational neuroscience, as the predicted patterns of dopaminergic activity have been seen in diverse studies in multiple species (Eshel, Tian, Bukwich, & Uchida, 2016; Tobler, Fiorillo, & Schultz, 2005; Zaghloul et al., 2009).

However, this classical theory does not account for the important role of dopamine in action planning. This role is evident from the difficulties in initiation of voluntary movements seen after the death of dopaminergic neurons in Parkinson’s disease. This role is consistent with the diversity in the activity of dopaminergic neurons, with many of them responding to movements (da Silva, Tecuapetla, Paixão, & Costa, 2018; Dodson et al., 2016; Howe & Dombeck, 2016; Jin & Costa, 2010; Lee, Mattar, Parker, Witten, & Daw, 2019; Schultz, Ruffieux, & Aebischer, 1983; Syed et al., 2016). The function of dopamine in energizing movements is likely to be underlined by the effects it has on the excitability or gain of the target neurons (Hernández-López et al., 2000; Thurley, Senn, & Luscher, 2008). Understanding the role of dopamine in action planning and movement initiation is important for refining treatments for Parkinson’s disease, where the symptoms are caused by dopamine depletion. Despite this importance, there is no mathematical framework, which can describe the role of dopamine in both learning and action planning.

A promising theory, called active inference, may provide the foundation for a framework accounting for such a dual role of dopamine (Friston, 2010). This theory relies of an assumption that the brain attempts to minimize prediction errors defined as the differences between observed stimuli and expectations. In active inference, these prediction errors can be minimized in two ways: through learning - by updating expectations to match stimuli, and through action - by changing the world to match the expectations. According to the active inference theory, prediction errors may need to be minimized by actions, because the brain maintains prior expectations that are necessary for survival and so cannot be overwritten by learning, e.g. food reserves should be at a certain level. When such predictions are not satisfied, the brain plans actions to reduce the corresponding prediction errors, e.g. by finding food.

This paper suggests that a more complete description of dopamine function can be gained by integrating reinforcement learning with elements of three more recent theories. First, taking inspiration from active inference, we propose that prediction errors represented by dopaminergic neurons are minimized by both learning and action planning, which gives rise to the roles of dopamine in both these processes. Second, we incorporate a recent theory of habit formation, which suggests that the habit and goal-directed systems learn on the basis of distinct prediction errors (Miller, Shenhav, & Ludvig, 2019), and we propose that these prediction errors are encoded by distinct populations of dopaminergic neurons, giving rise to the observed diversity of their responses. Third, we assume that the most appropriate actions are identified through Bayesian inference (Solway & Botvinick, 2012), and present a mathematical framework describing how this inference can be physically implemented in anatomically known networks within the basal ganglia. Since the framework extends the description of dopamine function to action planning, we refer to it as the DopAct framework. The DopAct framework accounts for a wide range of experimental data including the diversity of dopaminergic responses, the difficulties in initiation of voluntary movements under dopamine depletion, and it makes several experimentally testable predictions.

## Results

To provide an intuition for the DopAct framework, we start with giving its overview. Next, we formalize the framework, and then show examples of models developed within it for two tasks commonly used in experimental studies of reinforcement learning and habit formation: selection of action intensity (such as frequency of lever pressing) and choice between two actions.

### Overview of the framework

This section first gives an overview of computations taking place during action planning in the DopAct framework, and then summarizes how these computations could be implemented in neural circuits including dopaminergic neurons.

It has been proposed that the aim of action selection is to bring an animal to a desired level of reserves such as food, water, etc. (Hull, 1952; Stephan et al., 2016). Although there are multiple internal dimensions which animals need to optimize, e.g. temperature (Buckley, Kim, McGregor, & Seth, 2017), internal salt levels (Cone et al., 2016), etc., for simplicity, we will consider a single dimension of food reserves. As there exists an optimal level of food reserves, an animal should only seek food resources, if its reserves are below the optimum value. Here we propose, that the mechanisms, which the brain employs to achieve the desired reserves level, include a system that is able to compute how much resources should be acquired in a given situation, and a circuit that is able to select an action to obtain the desired reward. We refer to these two components as a valuation system and an actor, respectively. This paper focusses on describing the actor. Nevertheless, we briefly summarise the computations in the valuation system below, because it will help in understanding the computations of the actor.

In DopAct framework, the role of the valuation system is to compute how much resources the animal should aim at acquiring in a given situation. We distinguish between two classes of factors that describe a situation of an animal: internal factors discussed earlier to which we refer as ‘reserves’, and external factors related to the environment, such as stimulus or location in space, to which we refer as a ‘state’ following reinforcement learning terminology. Thus the role of the valuation system is to compute the value *ν* defined as the amount of resource, which should be acquired by the animal in a given state *s*. The value ν depends on both the amount of resources available in state *s*, and the current level of reserves, as illustrated in Figure 1A, where arrows denote animal’s estimate for the change in reserve levels that may be achieved in a given state. For example, imagine a whale in a state “by a big school of sardines”. If the reserves level resulting from consuming the entire resource available is still below the optimum level, as in the top display of Figure 1A, then the desired value *ν* is equal to the entire resource available (i.e. the whale should swallow the whole school). By contrast, the bottom display illustrates the case when consuming the whole resource would move the animal beyond the optimum reserves level, and here value *ν* is equal to the amount required to bring the food reserves to the desired level (i.e. the whale should only swallow a corresponding fraction of the school). To perform this computation, the valuation system needs to be able to learn how much resource is available in a given state (analogously to a critic in classical reinforcement learning), and during planning compute what fraction of that resource should be acquired.

**Figure 1.**
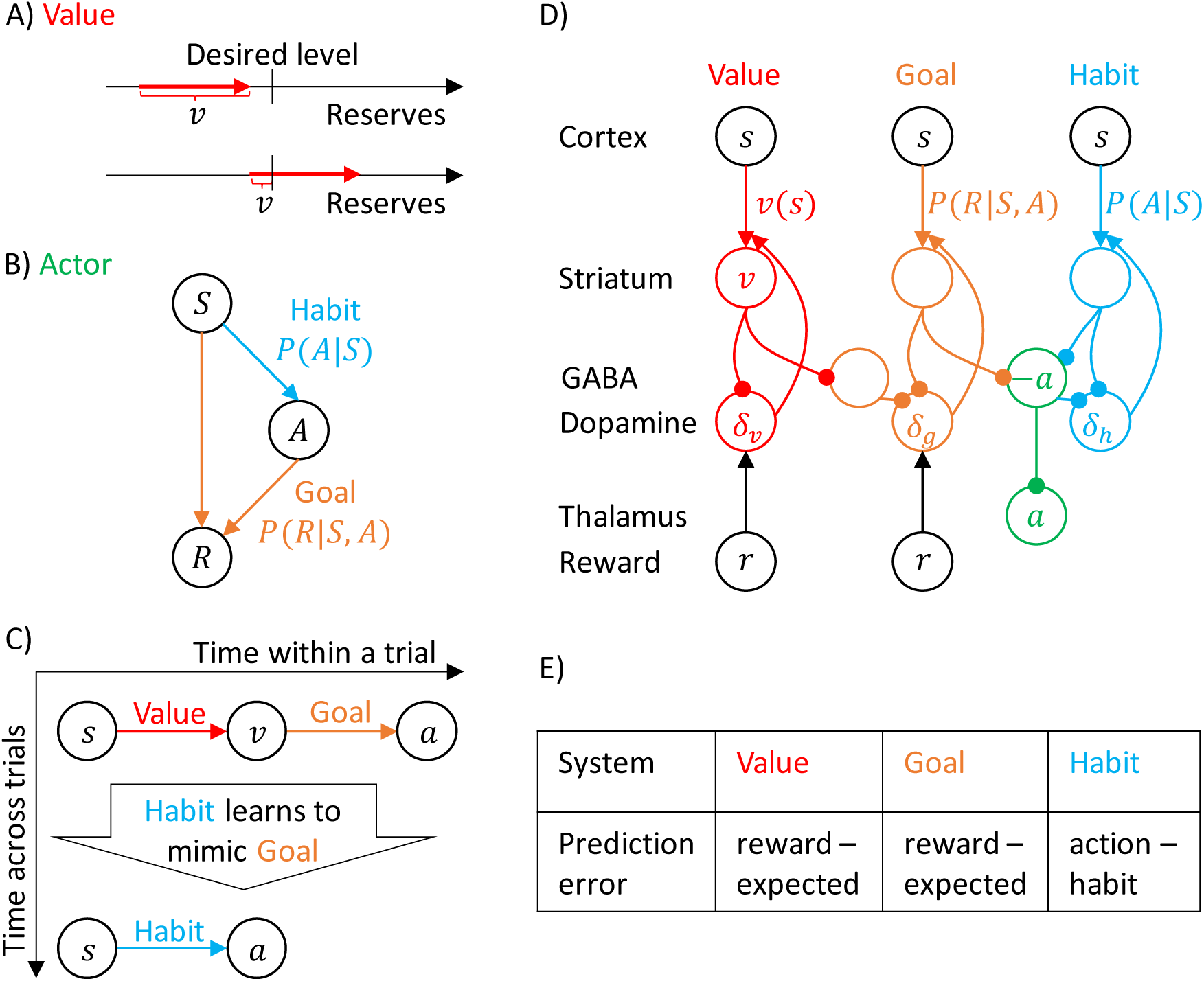
Overview of systems within the DopAct framework. A) Computation of the value for two different initial levels of reserves. Red arrows show changes in the reserves level that can be obtained in a current state. B) Probabilistic model learned by the actor. Random variables are indicated by circles, and arrows denote dependencies learned by different systems. C) Schematic overview of information processing in the framework at different stages of task acquisition. D) Mapping of the systems on different parts of the cortico-basal ganglia network. Circles correspond to neural populations located in the regions indicated by labels to the left, where ‘Striatum’ denotes medium spiny neurons expressing D1 receptors, ‘GABA’ denotes inhibitory neurons located in vicinity of dopaminergic neurons, and ‘Reward’ denotes neurons providing information on the magnitude of instantaneous reward. Arrows denote excitatory projections, while lines ending with circles denote inhibitory projections. E) Summary of prediction errors encoded in different systems.

Since this paper focusses on describing computations in the actor, for simplicity, we assume that the valuation system is able to compute the value *ν*, but this paper does not describe how that computation is performed. In simulations we mostly focus on a case of low reserves, where *ν* is equal to the whole resource available, and use a simple model similar to a critic in standard reinforcement learning, which just learns the average value *ν(s)* of resource in state *s* (Sutton & Barto, 1998), but does not consider reserve levels. Extending the description of the valuation system will be an important direction for future work and we come back to it in Discussion. In the considered case of low resources, for simplicity we use word ‘reward’ as a synonym of ‘resource’.

The goal of the actor is to select an action to obtain the reward set by the valuation system. This action is selected through inference in a probabilistic model, which describes relationships between states, actions and rewards. We denote random variables from which states, actions and rewards are sampled by *S, A* and *R*, and particular values of these variables by corresponding small letters. The DopAct framework assumes that two systems within the actor learn distinct relationships between the variables, shown in Figure 1B. The first system, shown in orange, learns how the reward depends on action selected in a given state, and we refer to it as ‘goal-directed’, because it can infer actions that typically lead to the desired reward. The second system, in blue, learns which actions should generally be chosen in a given state, and we refer to it as ‘habit’, because it suggests actions without considering the value of the reward currently available. Both goal-directed and habit systems propose an action, and their influence depends on their relative certainty.

Figure 1C gives an overview of how the systems mentioned above contribute to action planning, in a typical task. During initial trials of a task, the valuation system (shown in red) evaluates the current state *s* and computes the value of desired reward *ν*,and the goal-directed system selects the action *a*. At this stage the habit system contributes little to the planning process as its uncertainty is high. As the training progresses, the habit system learns to mimic the choices made by the goal-directed system (Miller et al., 2019). If for a given state *s*, the animal selects very similar actions over many trials, the certainty of the habit system increases. In this case, on later trials the action is mostly determined by the habit system (Figure 1C). Such choice is also faster, because it does not require an intermediate step of computing the value of the state.

The details of the above computations in the framework will be described in the next section, and it will be later shown how an algorithm inferring action can be implemented in a network resembling the anatomy of the basal ganglia (Figure 1D). But before going through a mathematical description, let us first provide an overview of this implementation. In this implementation, the valuation, goal-directed and habit systems are mapped on the spectrum of cortico-basal ganglia loops (Alexander, DeLong, & Strick, 1986), ranging from valuation in a loop including ventral striatum, to habit in a loop including the dorsolateral striatum that has been shown to be critical for habitual behaviour (Burton, Nakamura, & Roesch, 2015). In the DopAct framework, the probability distributions learned by the actor are encoded in the strengths of synaptic connections in the corresponding loops, primarily in cortico-striatal connections. As in a standard implementation of the critic (Houk et al., 1995), the parameters of the value function learned by the valuation system are encoded in cortico-striatal connections of the corresponding loop.

Analogous to classical reinforcement learning theory, dopaminergic neurons play a critical role in learning, and encode errors in predictions made by the systems in the DopAct framework. However, by contrast to the standard theory, dopaminergic neurons do not all encode the same signal, but instead dopaminergic populations in different systems compute errors in predictions made by their corresponding system (Figure 1E). Since both valuation and goal-directed systems learn to predict reward, the dopaminergic neurons in these systems encode reward prediction errors (which slightly differ between these two systems, as will be illustrated in simulations presented later). By contrast, the habit system learns to predict action on the basis of a state, so its prediction error encodes how the currently chosen action differs from a habitual action in the given state. Thus these dopaminergic neurons respond to non-habitual actions in the DopAct framework. We denote the prediction errors in the valuation, goal-directed and habit systems by *δ*_*ν*_, *δ*_*g*_ and *δ*_*h*_, respectively. In the DopAct framework, the dopaminergic neurons send these prediction errors to the striatum, where they trigger plasticity of cortico-striatal connections. For example, when an action is selected mostly by the goal-directed system, the prediction error in the habit system will trigger plasticity in the striatal neurons of the habit system, so they tend to predict this action in the future. In this way, the habit system learns to mimic the goal-directed system.

The systems communicate through an ‘ascending spiral’ structure of striato-dopaminergic projections identified by Haber, Fudge, and McFarland (2000). These Authors observed that dopaminergic neurons within a given loop project to the corresponding striatal neurons, while the striatal neurons project to the dopaminergic neurons in the corresponding and next loops, and they proposed that the projections to the next loop go via interneurons, so they are effectively excitatory (Figure 1D). In the DopAct framework, once the striatal neurons in the valuation system compute the value of the state *ν*, they communicate it to the goal-directed system via the dopaminergic neurons in the goal-directed system.

In the DopAct framework, dopamine in the goal-directed system plays a role in both action planning and learning, and now an overview of this role is given. In agreement with classical reinforcement learning theory, the dopaminergic activity *δ*_*g*_ encodes reward prediction error, namely the difference between the reward (including both obtained and available reward) and the expected reward (Schultz et al., 1997), but in the DopAct framework the expectation of reward only arises from formulating a plan to achieve it. Thus in the presence of reward, the prediction error *δ*_*g*_ can only be reduced to zero, once a plan to obtain the reward is formulated.

The dual role of dopamine in the DopAct framework stems from the two ways in which the goal-directed system minimizes reward prediction error: learning and action planning. To gain an intuition for how this system operates, let us consider a simple example of a naïve hungry rat exploring a conditioning apparatus. Assume that the rat presses a lever and a food pellet is delivered in a food port (Figure 2). The sight of this unexpected reward will trigger a dopamine response. According to the DopAct framework, the reward prediction error arises in the goal-directed system, because the valuation system noted that a reward is available, but the goal-directed system has not yet prepared actions to obtain the reward (so it has not formed an expectation). The resulting prediction error is being used in two ways. First, the prediction error triggers a process of planning actions that can get the reward. This facilitation of planning arises in the network, because the dopaminergic neurons in the goal-directed system project to striatal neurons (Figure 1D), and increase their excitability. Once an action plan has been formulated, the animal starts to expect the available reward, and the dopamine level encoding the prediction error decreases. Importantly, in this network dopamine provides a crucial feedback to striatal neurons on whether the formulated action plan is sufficient to obtain the available reward. If it is not, this feedback triggers changes in the action plan until it becomes appropriate. Thus the framework suggests why it is useful for neurons encoding reward prediction error to be involved in planning, namely it suggests that this prediction error provides a useful feedback for the action planning system, informing if the plan is suitable to obtain the reward.

**Figure 2.**
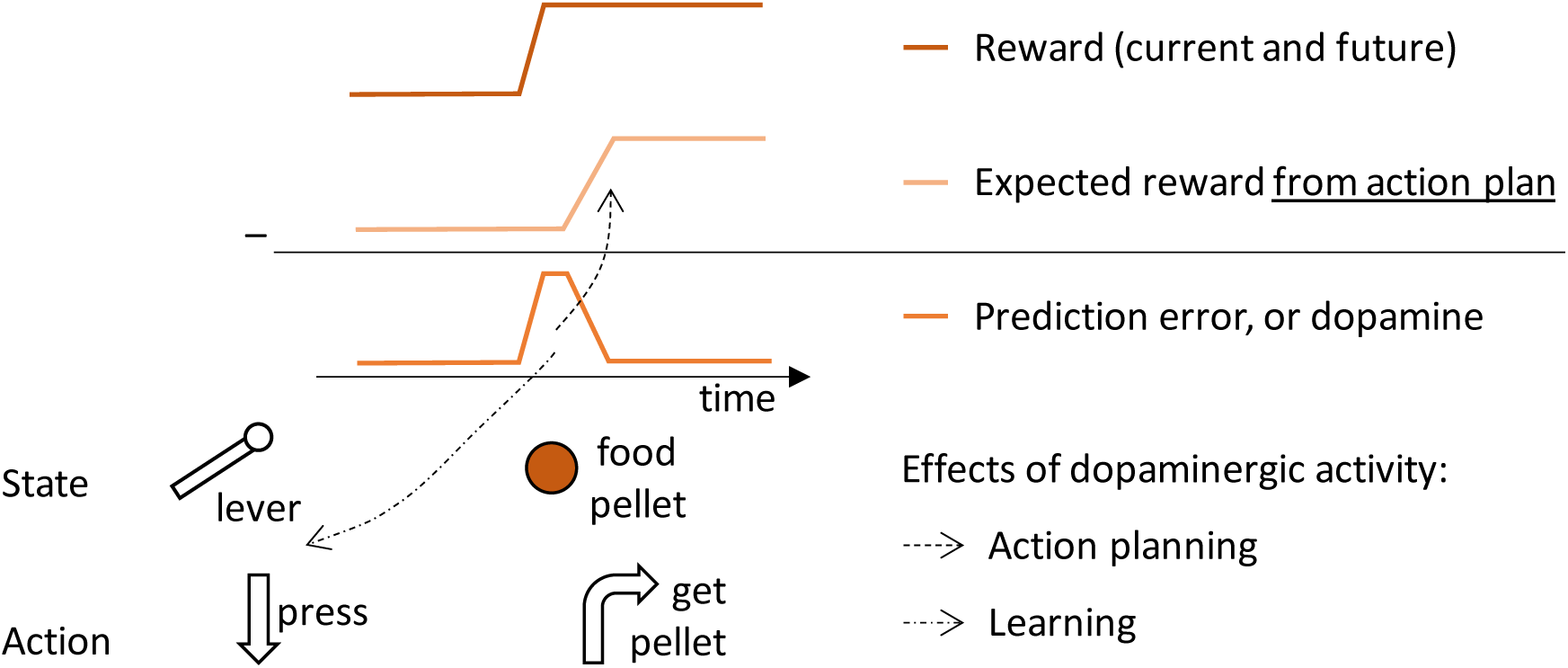
Schematic illustration of changes in dopaminergic activity in the goal-directed system while a naïve hungry rat presses a lever and a food pellet is delivered. The prediction error encoded in dopamine (bottom trace) is equal to a difference between the reward available (top trace) and the expectation of reward arising from a plan to obtain it (middle trace). The dashed arrows schematically indicate the processes such prediction error triggers.

Second, the prediction error allows the animal to learn that rewards are available after certain actions at particular states, so in this case, it will modify synaptic connections encoding the value of lever pressing.

### Formal description of the framework

Let us now provide the details of the DopAct framework. For clarity, we will follow Marr’s levels of description, and in this section, we discuss computation and algorithm employed by the actor, while sample implementations for two commonly used tasks are presented in the following sections. To illustrate the computations in the framework we will consider a simple task, in which only an intensity of a single action needs to be chosen. Such choice needs to be made by animals in classical experiments investigating habit formation, where the animals are offered a single lever, and need to decide how frequently to press it. Furthermore, action intensity often needs to be chosen by animals also in the wild (e.g. a cat deciding how vigorously to pounce on a prey, a chimpanzee choosing how strongly to hit a nut with a stone, or a sheep selecting how vigorously to eat the grass). Let us denote the action intensity by *a*. Let us assume that the animal chooses it on the basis of the reward it expects *R* and the stimulus *s* (e.g. size of prey, nut or height of grass). Thus the animal needs to infer an action intensity sufficient to obtain the desired reward (but not larger to avoid unnecessary effort).

Let us first describe the computations performed in the DopAct framework. As mentioned in the previous section, we assume that the actor maintains two probability distributions: The goal-directed system encodes how the reward depends on states and actions, while the habit system encodes the probability distribution of generally selecting actions in particular states. During action planning, when an animal notices reward available *R* = *ν*, it combines information from both systems through Bayesian inference. According to Bayes’ theorem (Equation 3.1 in Figure 3), the posterior probability of selecting a particular action given available reward is proportional to the product of a likelihood of the reward given the action, which we propose is represented in the goal-directed system, and a prior, which we propose is encoded by the habit system.

**Figure 3.**
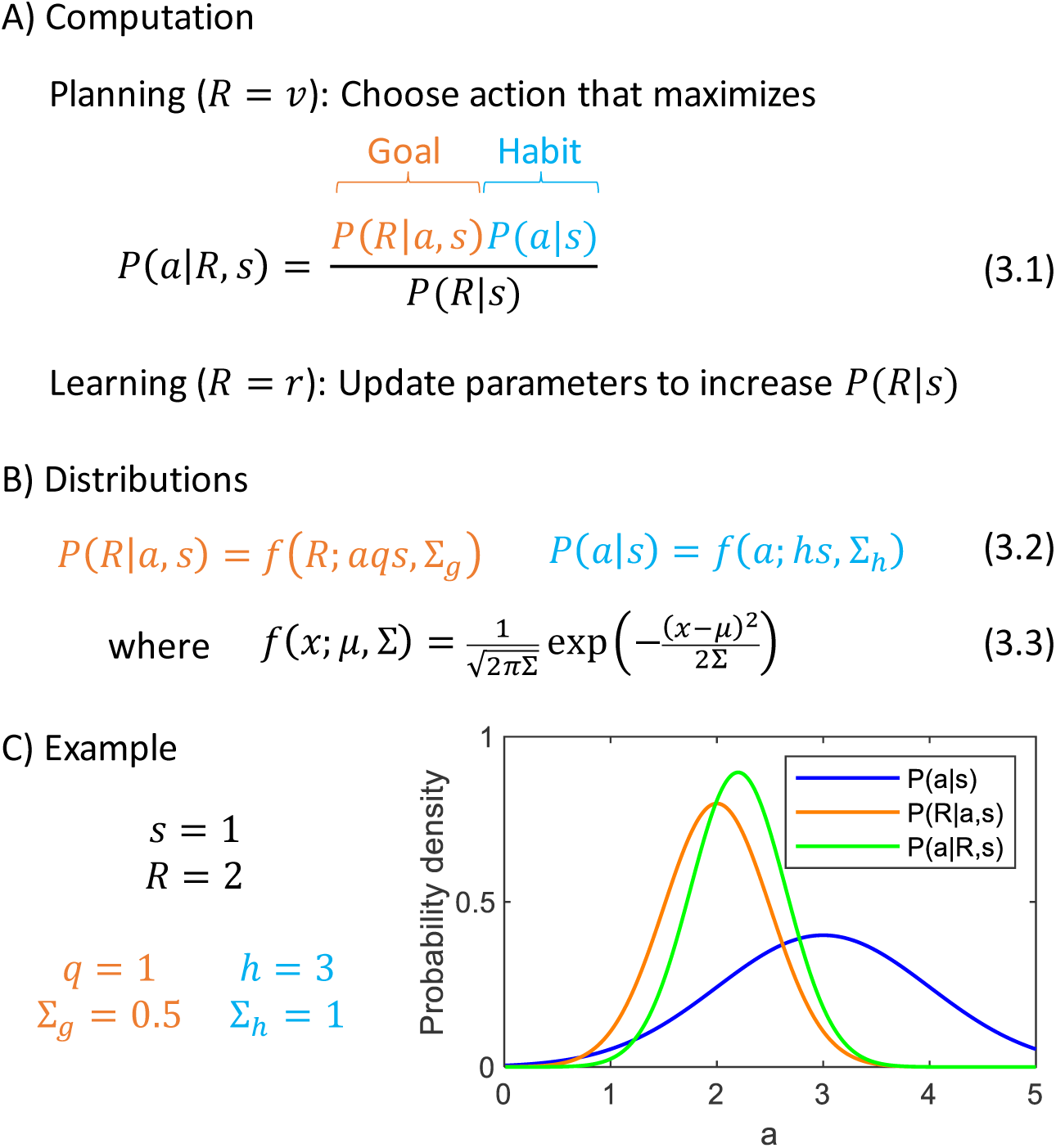
Computational level. (A) Summary of computations used by the actor in the DopAct framework. (B) Sample form of probability distributions. (C) An example of inference of action intensity. In this example the stimulus intensity is equal to *s* = 1, the valuation system computes desired resource *R* = 2, and the parameters of the probability distributions encoded in the goal-directed and habit systems are listed in the panel. The blue curve shows the distribution of action intensity, which the habit system has learned to be generally suitable for this stimulus. The orange curve shows probability density of obtaining reward of 2 for a given action intensity, and this probability is estimated by the goal-directed system. For the chosen parameters, it is the probability of obtaining 2 from a normal distribution with mean *a*. Finally, the green curve shows a posterior distribution computed from Equation 3.1.

In the DopAct framework, an action *a* is selected which maximizes the probability *P(a|R,s)*. An analogous way of selecting actions has been used in models treating planning as inference (Attias, 2003), and it has been nicely summarized by Solway and Botvinick (2012): “The decision process takes the occurrence of reward as a premise, and leverages the generative model to determine which course of action best explains the observation of reward.” In this paper, we make explicit the rationale for this approach: The desired amount of resources that should be acquired depends on the levels of reserves (and a given state); this value is computed by the valuation system, and the actor needs to find the action depending on this reward. Let us provide a further rationale for selecting an action *a* which maximizes *P(a| R, s)*, by analysing what this probability expresses. Let us consider a following hypothetical scenario: An animal selected an action without considering the desired reward, i.e. by sampling it from its default policy *P(a|s)* provided by the habit system, and obtained reward *R*. In this case, *P(a|R, s)* is the probability that the selected action was *a*. When an animal knows the amount of resource desired *R*, then instead of just relying on the prior, the animal should rather choose an action maximizing *P(a|R,s)*, which was the action most likely to yield this reward in the above scenario.

One may ask why it is useful to employ the habit system, instead of exclusively relying on the goal-directed system that encodes the relationship between rewards and actions. The answer is that there may be uncertainty in the action suggested by the goal-directed system, arising for example, from noise in the computations of the valuation system or inaccurate estimates of the parameters of the goal-directed system. According to Bayesian philosophy, in face of such uncertainty, it is useful to additionally bias the action by a prior, which here is provided by the habit system. This prior encodes an action policy that has overall worked in the situations previously experienced by the animal, so it is a useful policy to consider under uncertainty in the goal-directed system. Later we will show in simulations that incorporating the prior may indeed help to select more optimal actions.

To provide an intuition for how the action intensity is computed, let us consider an example. Let us specify the form of the prior and likelihood distributions. They are given in Figure 3B, where *f (x; μ, Σ)* denotes the probability density of a normal distribution with mean *μ* and variance Σ. In a case of the prior, we assume that action intensity is normally distributed around a mean given by stimulus intensity scaled by parameter *h*, reflecting an assumption that a typical action intensity often depends on a stimulus (e.g. the larger a nut, the harder a chimpanzee must hit it). On the other hand, in a case of the probability of reward *R* maintained by the goal-directed system, the mean of the reward is equal to a product of action intensity and the stimulus size, scaled by parameter *q.* We assume that the mean reward depends on a product of *a* and s for three reasons. First, in many situations reward depends jointly on the size of the stimulus, and the intensity with which the action is taken, because if the action is too weak, the reward may not be obtained (e.g. a prey may escape or a nut may not crack), and the product captures this dependence of reward on a conjunction of stimulus and action. Second, in many foraging situations, the reward that can be obtained within a period of time is proportional to a product of *a* and s (e.g. amount of grass eaten by a sheep is proportional to both how vigorously the sheep eats it, and how high the grass is). Third, when the framework is generalized to multiple actions later in the paper, the assumption of reward being proportional to a product of *a* and s will highlight a link with classical reinforcement learning. We denote the variances of the distributions of the goal-directed and habit systems by Σ_*g*_ and Σ_*h*_. Figure 3C shows an example of probability distributions encoded by the two systems for sample parameters. It also shows a posterior distribution *P(a|R, s)*, and please note that its peak is in between the peaks of the distributions of the two systems, but it is closer to the peak of a system with smaller uncertainty (orange distribution is narrower). This illustrates how in the DopAct framework, the action is inferred by incorporating information from both systems, but weighting it by the certainty of the systems.

In addition to action planning, the animal needs to learn from the outcomes, to predict rewards more accurately in the future. After observing the reward actually obtained *R* = *r*, the parameters of the distributions should be updated to increase *P(R|s)*, so in the future the animal is less surprised by the reward obtained in that state (Figure 3A).

Let us now describe an algorithm used by the actor to infer action intensity *a* that maximizes the posterior probability *P(a|R,s)*. This posterior probability can be computed from Equation 3.1, but note that *a* does not occur in the denominator of that equation, so we can simply find the action that maximizes the numerator. Hence, we define an objective function *F* equal to a logarithm of the numerator of Bayes’ theorem (Equation 4.1 in Figure 4). Introducing the logarithm will simplify function *F* because it will cancel with exponents present in the definition of normal density (Equation 3.3), and it does not change the position of the maximum of the numerator because the logarithm is a monotonic function. For example, the green curve in Figure 4B shows function *F* corresponding to the posterior probability in Figure 3C. Both green curves have the maximum at the same point, so instead of searching for a maximum of a posterior probability, we can seek the maximum of a simpler function *F*.

**Figure 4.**
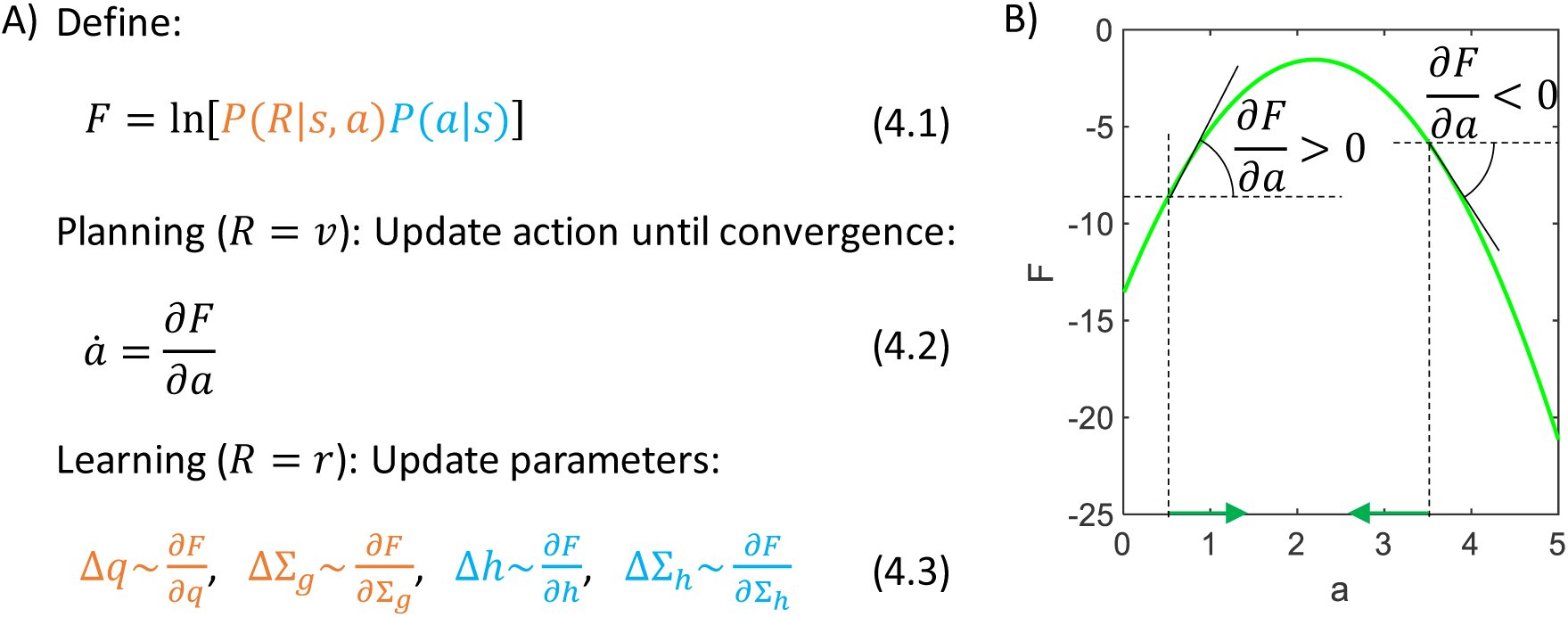
Algorithmic level. (A) Summary of the algorithm used by the actor. (B) Identifying an action based on a gradient of *F*. The panel shows an example of a dependence of *F* on *a*, and we wish *a* to take the value maximizing *F*. To find the action, we let *a* to change over time in proportion to the gradient of *F* over *a* (Equation 4.2, where the dot over *a* denotes derivative over time). For example, if the action is initialized to *a* = 1.5, then the gradient of *F* at this point is positive, so *a* is increased (Equation 4.2), as indicated by a green arrow on the x-axis. These changes in *a* continue until the gradient is no longer positive, i.e. when *a* is at the maximum. Analogously, if the action is initialized to *a* = 3.5, then the gradient of *F* is negative, so *a* is decreased until it reaches the maximum of *F*.

During action planning we set reward to reward available *R* = *ν* in Equation 4.1, and we find the action maximizing *F*. This can be achieved by initializing *a* to any value, and then changing it proportionally to the gradient of *F* (Equation 4.2). Figure 4B illustrates that with such dynamics, the value of *a* approaches a maximum of *F*. Once *a* converges, the animal may select the action with the corresponding intensity. In summary, this method yields a differential equation describing an evolution of a variable *a*, which converges to a value of *a* that maximizes *P(a|R, s)*.

After obtaining a reward, *R* is set to the reward obtained *R* = *r* in Equation 4.1, and the values of parameters are changed proportionally to the gradient of *F* (Equations 4.3). Such parameter updates allow the model to be less surprised by the rewards (as aimed for in Figure 3A), because under certain assumptions function *F* expresses “negative free energy”, which provides a lower bound on *P(R|s)* (Friston, 2005) (a detailed explanation for why *F* expresses negative free energy for an analogous problem is given by Bogacz (2017)). Thus changing the parameters to increase *F*, rises the lower bound on *P (R|s)*, and so it tends to increase *P(R|s)*.

### Network selecting action intensity

This section describes how the above algorithm for inferring action intensity could be implemented in a network resembling basal ganglia anatomy. Let us start with considering a special case in which both variance parameters are fixed to Σ_*g*_ = Σ_*h*_ = 1, because then the mapping of the algorithm on the network is particularly beautiful.

Let us derive the details of the algorithm (general form of which is given in Figure 4A) for the problem choosing action intensity. Substituting probability densities of likelihood and prior distributions (Equations 3.2-3.3) for the case of unit variances into Equation 4.1 (and ignoring constants 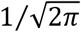), we obtain the expression for the objective function *F* in Equation 5.1. We see that *F* consists of two terms, which are the squared prediction errors associated with goal-directed and habit systems. The prediction error for the goal-directed system describes how the reward differs from the expected mean, while the prediction error of the habit system expresses how the chosen action differs from that typically chosen in the current state (Equations 5.2). Equation 5.1 highlights that inferring action intensity that maximizes *F* corresponds to reducing prediction errors. As described in the previous section, action intensity can be found by changing its value according to a gradient of *F* (Equation 4.2). Computing the derivative of *F* over *a*, we obtain Equation 5.3, where the two colours indicate terms connected with derivatives of the corresponding prediction errors. Finally, when the reward is obtained, we modify the parameters proportionally to the derivatives of *F* over the parameters, which are equal to relatively simple expressions in Equations 5.4.

The key features of the algorithm in Figure 5A naturally map on the known anatomy of striato-dopaminergic connections. This mapping relies on three assumptions analogous to those typically made in models of the basal ganglia: (i) the information about state s is provided to the striatum by cortical input, (ii) the parameters of the systems *q* and *h* are encoded in the cortico-striatal weights, and (iii) the computed action intensity is represented in the thalamus (Figure 5B). Under these assumptions, Equation 5.3 describing an update of action intensity can be mapped on the circuit: The action intensity in the model is jointly determined by the striatal neurons in the goal-directed and habit systems, which compute the corresponding terms of Equation 5.3, and communicate them by projecting to the thalamus via the output nuclei of the basal ganglia. The first term *δ*_*g*_*qs* can be provided by striatal neurons in the goal-directed system (denoted by *G* in Figure 5B): They receive cortical input encoding stimulus intensity *s*, which is scaled by cortico-striatal weights encoding parameter q, so these neurons receive synaptic input *qs*. To compute *δ*_*g*_*qs*, the gain of the striatal neurons in the goal-directed system needs to be modulated by dopaminergic neurons encoding prediction error *δ*_*g*_ (this modulation is represented in Figure 5B by an arrow from dopaminergic to striatal neurons). Hence, these dopaminergic neurons drive an increase in action intensity until the prediction error they represent is reduced (as discussed in Figure 2). The second term *hs* in Equation 5.3 can be computed by a population of neurons in the habit system receiving cortical input via connection with the weight *h*. Finally, the last term —*a* simply corresponds to a decay. Moreover, according to Equations 5.4, the prediction errors modulate the plasticity of cortico-striatal connections in both systems (represented in Figure 5B by arrows going from dopamine neurons to parameters).

**Figure 5.**
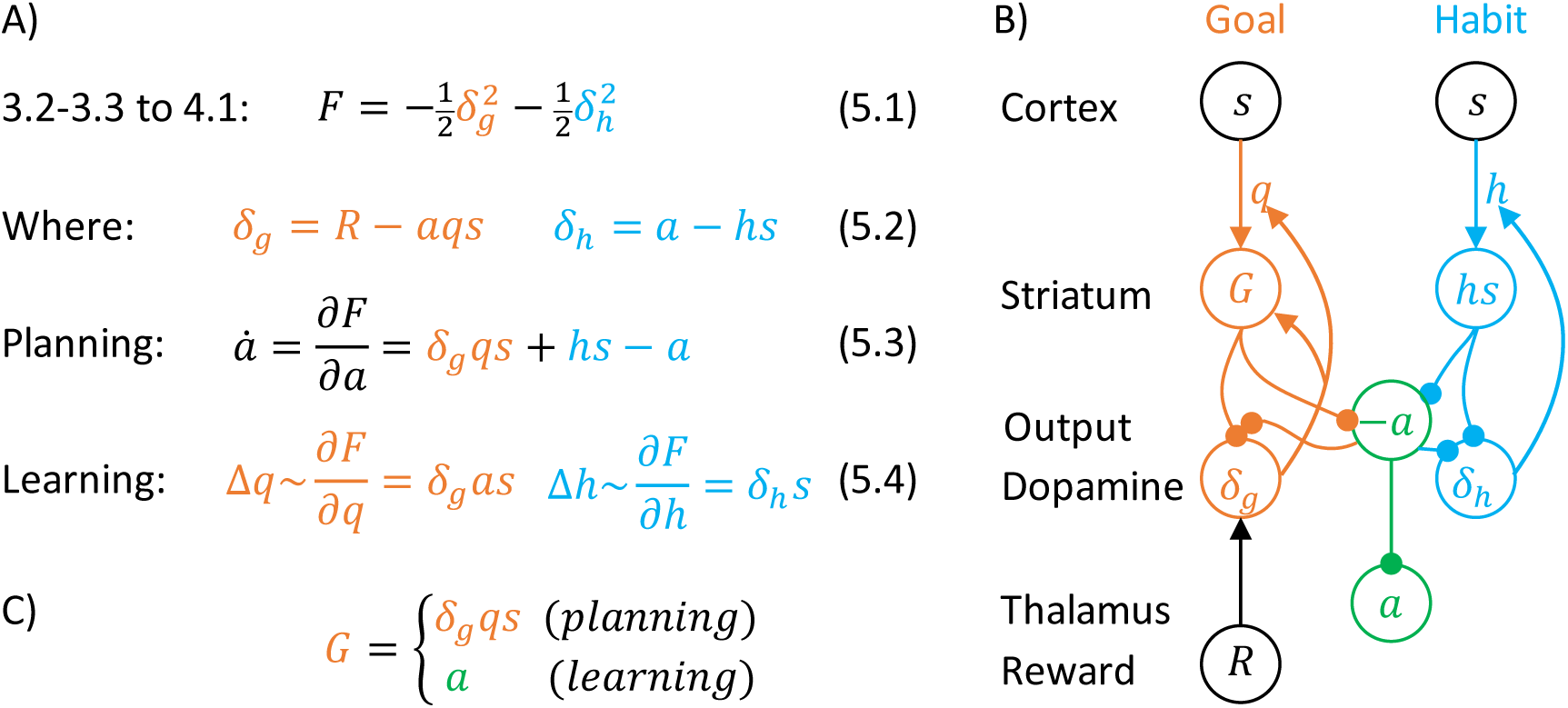
Description of a model selecting action intensity, in a case of unit variances. A) Details of the algorithm employed in the model. B) Mapping of the algorithm on network architecture. Notation as in Figure 1D, and additionally “Output” denotes the output nuclei of the basal ganglia. C) Definition of striatal activity in the goal-directed system.

There are several ways of mapping the remaining details of the algorithm on the striato-dopaminergic circuit, and as a proof of principle we show here one such mapping, as an example. Within each system, dopaminergic neurons compute errors in the predictions about the corresponding variable, i.e. reward for the goal-directed system, and action for the habit system. In the habit system, the prediction error is equal to a difference between action *a* and expectation *hs* (blue Equation 5.2). Such error can be easily computed in a network of Figure 5B, where the dopaminergic neurons in the habit system receive effective input form the output nuclei equal to *a* (as they receive inhibition equal to —*a*), and inhibition *hs* from the striatal neurons. In the goal-directed system, the prediction error is proportional to the difference between the reward and the expectation (orange Equation 5.2). The neurons computing prediction error in the goal-directed system in the network in Figure 5B receive input equal to reward, so let us now consider where the inhibition equal to the expectation *aqs* could originate. The prediction error node receives a related term *δ*_*g*_*qs* from the striatum (Figure 5B), and this input could be normalized by the activity of the error node itself *δ*_*g*_, to result in an effective input *qs*. Such input would still need to be scaled by action intensity *a* to compute the expectation. Further work is required to understand how such scaling may be achieved and one possibility is thorough an input from the output nuclei (included in Figure 5B) (Watabe-Uchida, Zhu, Ogawa, Vamanrao, & Uchida, 2012).

The prediction errors are used to update the parameters of the distributions represented by the systems, which are encoded in the weights of cortico-striatal connections. Once the actual reward is obtained, the learning of these parameters could be achieved through local synaptic plasticity dependent on dopaminergic modulation. In the goal-directed system, orange Equation 5.4 corresponds to local plasticity, if at the time of reward the striatal neurons encode information about action intensity (see definition of *G* in Figure 5C). Such information could be provided from the thalamus during action execution. Then the update of synaptic weight encoding parameter *q* will correspond to a standard three-factor rule (Kuśmierz, Isomura, & Toyoizumi, 2017) involving a product of presynaptic *(s)*, postsynaptic (a) and dopaminergic activity (*δ*_*g*_). The update of a weight encoding parameter *h* (blue Equation 5.4) is simply proportional to the product of presynaptic *(s)* and dopaminergic activity (*δ*_*h*_) for the simple model described in this section.

The Methods section shows how the model can be extended so that the parameters Σ_*g*_ and Σ_*h*_ describing variances of distributions are encoded in synaptic connections or internal properties of the neurons (such as leak conductance). In such an extended model, the action proposals of the two systems are weighted according to their certainties. As described in the Methods, a simple model of the valuation system based on a critic from standard reinforcement learning is employed in simulations (because the simulations correspond to a case of low level of animal’s reserves). Striatal neurons in the valuation system compute the reward expected in a current state as *ν* = *ws*, where *w* is a parameter encoded in cortico-striatal weights. The dopaminergic neurons in the valuation system encode the prediction error similar to that in the temporal-difference learning model, and after reward delivery, they modulate plasticity of cortico-striatal connections. The Method section also provides details of the implementation and simulations of the model.

### Simulations of action intensity selection

To illustrate how the model mechanistically operates and to help relate it to experimental data, we now describe a simulation of the model inferring action intensity. On each simulated trial the model selected action intensity, after observing a stimulus, which was set to *s* = 1. The reward obtained depended on action intensity as shown in Figure 6A, according to *r* = 5tanh(*a*/5) — 0.1*a*. Thus, the reward was proportional to the action intensity, transformed through a saturating function, and a cost was subtracted proportional to the action intensity, that could correspond to a price for making an effort. We also added small Gaussian noise to reward with standard deviation *σ*_*r*_ = 0.5 (to account for randomness in the environment), and to action intensity with standard deviation *σ*_*α*_ = 0.5 (to account for imprecision of the motor system or exploration).

**Figure 6.**
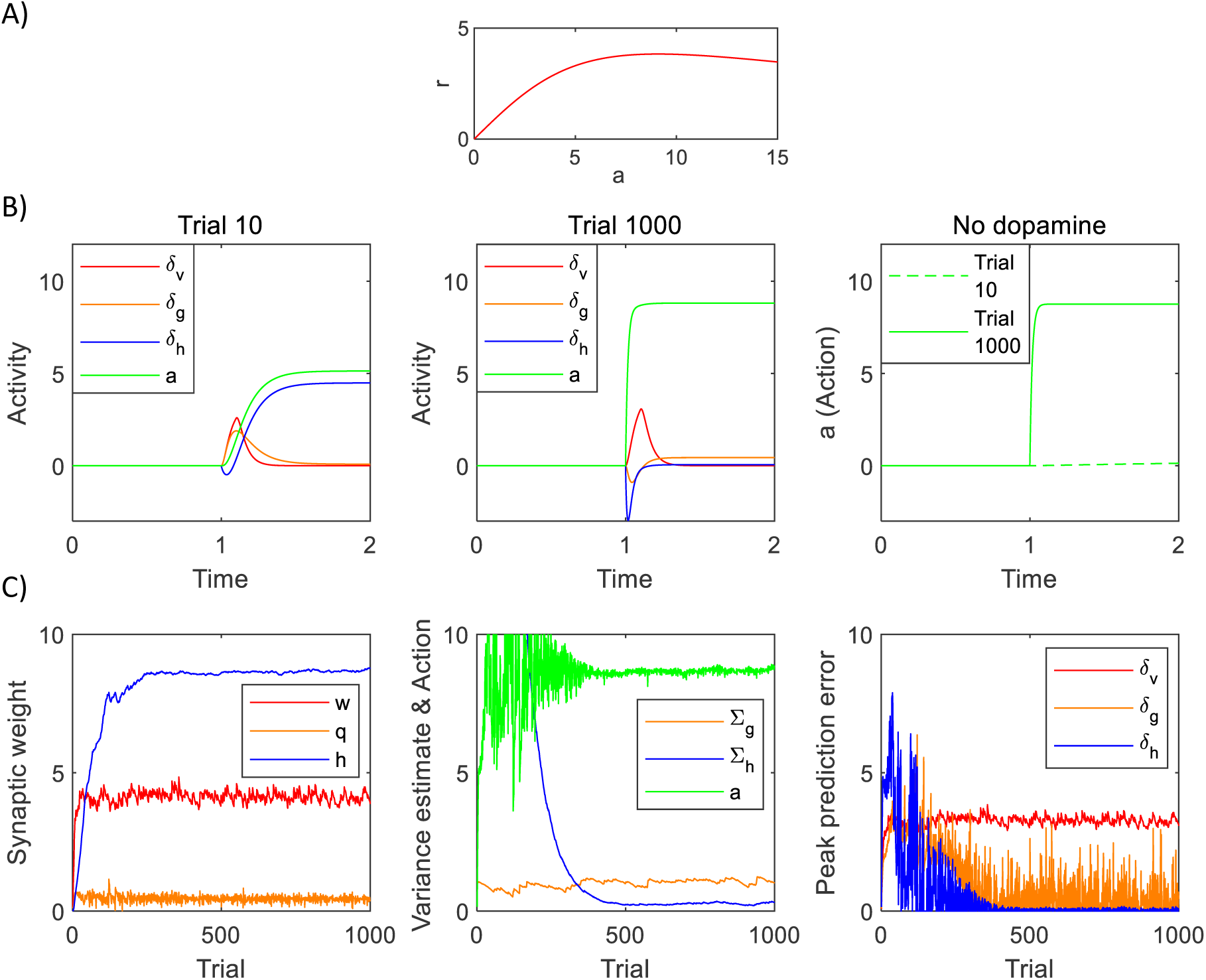
Simulation of a model selecting action intensity. A) Mean reward given to a simulated agent as a function of action intensity. B) Changes in model variables within a trial of a simulation. C) Changes in model variables and parameters across trials. The green curve in the middle display shows the action intensity at the end of a planning phase of each trial. The right display shows the maximum values of the prediction errors over the entire planning phase of each trial.

Figure 6B shows how the prediction errors and action intensity changed within a trial. The left display corresponds to one of the early trials. The stimulus was presented at time 1. The valuation system detected that a reward was available, which initially resulted in a prediction error in the goal-directed system, visible as an increase in the orange curve. This prediction error triggered a process of action planning, so with time the green curve representing planned action intensity increased. Once the action plan has been formulated, it provided a reward expectation, so the orange prediction error decreased. As the habit system has not formed significant habits on early trials, it was surprised by the chosen action, and this high value of blue prediction error drove its learning over trials. For simplicity, the simulation is shown for the entire period of 2 time units, but in a real neural system, the action is likely to be executed once *a* converges, and the resulting movements would change *a* and thus the habit prediction error. Therefore, the increase in the habit prediction error would be transient rather than sustained (as depicted in the figure). Extending the model to detect convergence of action intensity will be an important direction of future work and we come back to it in the Discussion.

The middle display in Figure 6B shows the same quantities on a trial that followed an extensive training. Now the habit system was highly trained and rapidly drove action planning, so the green curve showing planned action intensity increased more rapidly and to a higher level. Nevertheless, due to the dynamics in the model, the increase in action intensity was not instant, so there was a transient negative prediction error in the habit system while an action was not yet equal to the value predicted by the habit system. After this transient, the habit system was no longer surprised by the planned action, thus the blue prediction error converged to a value close to 0. At this stage of training, the goal-directed system was too slow to lead action planning, so the orange prediction error was lower. Finally, the red prediction error in the valuation system behaved as expected from the standard temporal difference learning model, i.e. a response to a stimulus predicting a reward was produced even after extensive training.

Dopaminergic neurons in the model are only required to facilitate planning in the goal-directed system, where they increase excitability of striatal neurons, but not in the habit system. To illustrate it, right display in Figure 6B shows simulations of a complete dopamine depletion in the model. It shows action intensity produced by the model in which following training, all dopaminergic neurons were set to 0. After just 9 trials of training, on the 10th trial, the model was unable to plan an action. By contrast, after 999 training trials, the model was still able to produce a habitual response, because dopaminergic neurons are not required for generating habitual responses in the model. This parallels the experimentally observed robustness of habitual responses to blocking dopaminergic modulation (Choi, Balsam, & Horvitz, 2005).

Figure 6C shows how the quantities in the model evolved over the trials in the simulation. The left display shows how the parameters in the three systems changed during learning. The parameter of the valuation system correctly converged to the maximum value of the reward available in the task *w* ≈ 4 (i.e. the maximum of the curve in Figure 6A). The parameter of the habit system correctly converged to action intensity resulting in the maximum reward *h* ≈ 9 (Figure 6A). The parameter of the goal-directed system converged to a vicinity of *q* ≈ 4/9, which allows the goal-directed system to expect the reward of 4 after selecting an action with intensity 9 (see orange Equation 3.2).

The middle display in Figure 6C shows how the variance parameters in the goal-directed and habit systems changed during the simulation. The variance of the habit system was initialised to a high value, and it decreased over time, resulting in an increased confidence of the habit system. The middle display also shows the action intensity produced in the model. On average it was close to *a* ≈ 9, which was approximately the value resulting in the maximum reward (Figure 6A). Although the model discovered this optimal action intensity quite rapidly, on a first few hundred trials, the inferred value of action was quite variable, because it relied on the inference in the goal-directed system, which in turn relied on the valuation system. The fluctuations in the parameters of these systems arising from learning from noisy rewards resulted in variable action intensity. However, once the habit system became more confident, the produced action intensity became closer to the optimum value, as the fluctuations in its parameter *h* were lower than for the other system (left display in Figure 6C). This illustrates the benefits of Bayesian inference mentioned earlier, that in the face of imprecision in the goal-directed system, a more accurate action intensity can be identified by also relying on a prior, which here was encoded in the habit system.

The right display in Figure 6C shows prediction errors after the stimulus in the three systems. The prediction error of the valuation system simply followed the value of the stimulus estimated by that system. The prediction error in the goal-directed system decreased on later trials, when it became slower than the habit system and no longer could lead action planning. There was a prediction error in the habit system on the initial trials, but it decreased once the action became habitual.

Although we demonstrated that including the habit system brings benefits of more accurate actions (Figure 6C, middle) and faster planning (Figure 6B), it may also have costs of excessive perseveration. Such perseveration is typically studied experimentally in tasks where rats have to press a lever multiple times to get a reward. As mentioned earlier, such tasks could be conceptualized as a choice of the frequency of pressing a lever, that could also be described by a single number *a*. Furthermore, the average reward rate experienced by an animal may correspond to a non-monotonic function similar to that in Figure 6A, because with more presses more rewards is typically received, but beyond a certain frequency, there may be a large effort cost. Therefore, we will continue to use the model described above to illustrate key qualitative effects known in the literature.

Figure 7A shows a simulation paralleling the reward omission protocol. The model has been simulated with reward depending on action intensity as in Figure 6A for a particular number of trials shown on x-axis of Figure 7A, and then the reward was not delivered so in simulations *r* = −0.1*a* reflecting just a cost connected with making an effort. The figure presents the average action frequency selected by the model after 500 trials since the rewards had been omitted. The action frequency increases with longer training, which parallels experimental observation of decreased sensitivity to reward omission with increased training (Dickinson, Squire, Varga, & Smith, 1998). When the rewards become omitted, the optimal action frequency is *a* = 0, as there is no benefit for action, but only a cost. Nevertheless, if the model had made the same responses for many trials, the habit system increased its confidence to the extent that it continued to drive action planning. In that case, the chosen action was close to the habitual one, so there was little prediction error in the habit system (*δ*_*h*_ ≈ 0), despite no reward, hence the habit system did not adjust significantly its behaviour or even its confidence. Although such perseveration in absence of rewards is not useful for animals, it is a cost paid in this neural circuit for the important benefits of fast and accurate habitual actions.

**Figure 7.**
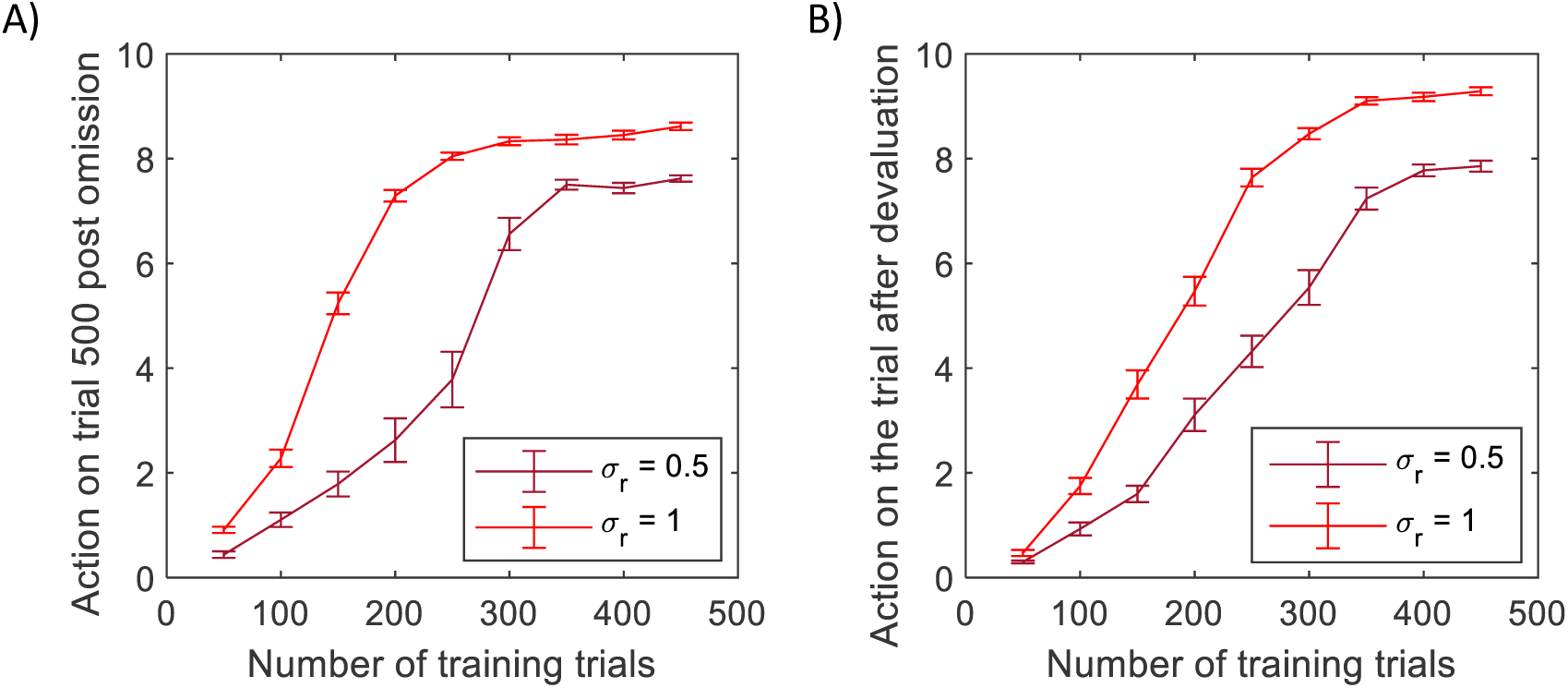
Produced action frequency following a change in task contingency or physiological state. A) Simulation of the omission paradigm. B) Simulation of the devaluation paradigm. Simulations were repeated 25 times, and error bars indicate standard error.

Figure 7B shows simulations of the model in a devaluation paradigm. In this paradigm, animals are trained to press a level for reward (e.g. food), and following this training the reward is devalued (e.g. the animals are fed to satiety, so they no longer desire the reward). It has been observed that animals trained for a short period would not press a lever following devaluation, while after an extensive training, the animal would press the lever, even though they no longer desire the reward (Dickinson, 1985). To simulate this paradigm, the model was trained for a number of trials (shown on x-axis), and then to simulate devaluation, the value computed by the valuation system was set to *ν* = 0 throughout the subsequent trial, as in a previous modelling study (Solway & Botvinick, 2012). Figure 7B shows the average action frequency inferred on this trial. It increased with training, analogously as in the devaluation experiments (Dickinson, 1985).

Following a previous modelling study of habit formation (Miller et al., 2019), Figure 7 also shows that the perseveration is more likely when the rewards are noisier. This effect occurs, because increasing noise in reward increases the estimate of reward variance Σ_*g*_, and this parameter also encodes the uncertainty of the goal-directed system. The increased uncertainty of the goal-directed systems makes it more likely to give in to the habit system. This property of the model parallels experimental observation that habitual behaviour is easier to produce in an experimental paradigm (variable interval schedule) (Dickinson, Nicholas, & Adams, 1983), which has more variable reward probability (Miller et al., 2019).

In summary, this section described a model for a simple task in which an animal had to select an intensity of a single action. Simulations of the model revealed a rich pattern of responses of different populations of dopaminergic neurons at different stages of task acquisition, and we will relate them to experimental data in Discussion.

### Choice between two actions

This section shows how models developed within the DopAct framework can also describe more complex tasks with multiple actions and multiple dimensions of state. We consider a task of choice between two options, often used in experimental studies, as it allows illustrating the generalization, and at the same time results in a relatively simple model. This section will also show that the models developed in the framework can under certain assumptions be closely related to previously proposed models of reinforcement learning and habit formation.

To make dimensionality of all variables and parameters explicit, we will denote vectors with a bar and matrices with a bold font. Thus 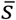 is a vector where different entries correspond to intensities of different stimuli in an environment, and 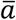 is a vector where different entries correspond to intensities of different actions. Equation 8.1 in Figure 8 shows how the definitions of the probability distributions encoded by the goal-directed and habit systems can be generalized to multiple dimensions. Orange Equation 8.1 states that the reward expected by the goal-directed system has mean 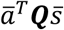, where ***Q*** is now a matrix of parameters. This notation highlights the link with the standard reinforcement learning, where the expected reward for selecting action *i* in state *j* is denoted by *Q*_*i,j*_: Note that if 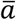 and 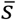 are both binary vectors with entries *i* and *j* equal to 1 in the corresponding vectors, and all other entries equal to 0, then 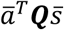 is equal to the element *Q*_*i,j*_ of matrix ***Q***.

**Figure 8.**
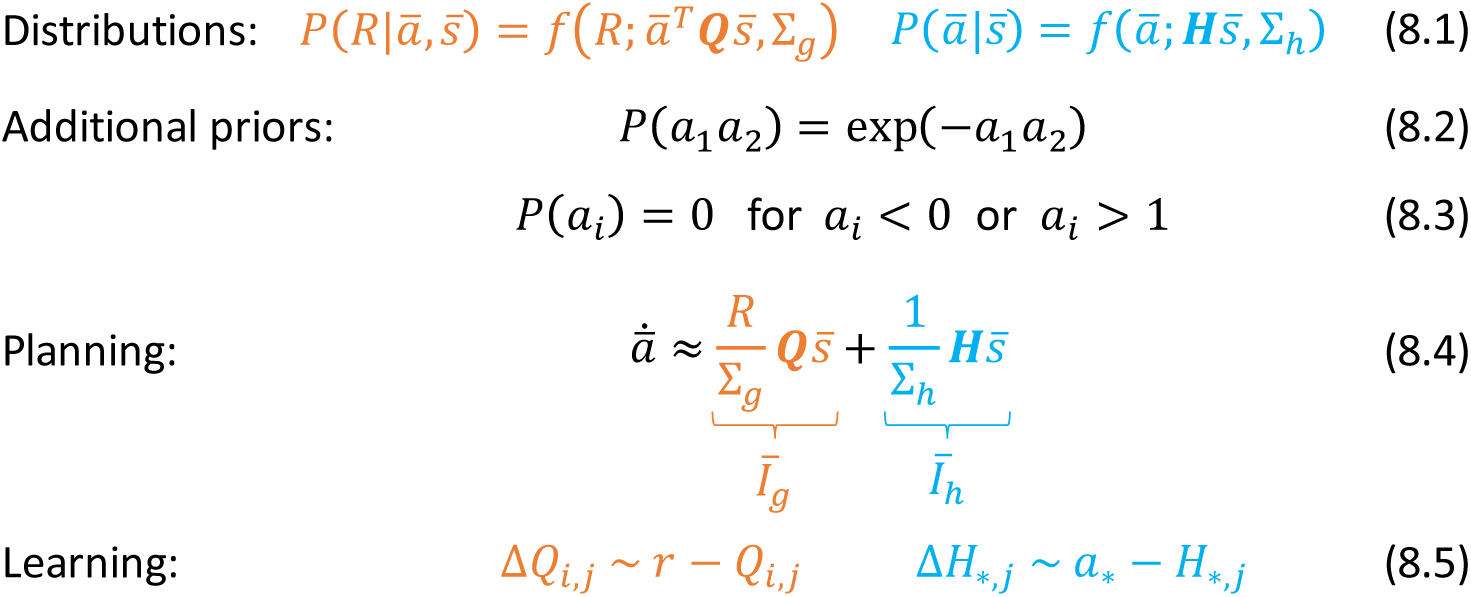
Summary of mathematical description of the model of choice between two options. In blue Equation 8.5, * indicates that the parameters are updated for all actions.

In the model, the prior probability is proportional to a product of three distributions. The first of them is encoded by the habit system and given in blue Equation 8.1. The expected action intensity encoded in the habit system has mean 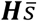, and this notation highlights the analogy with a recent model of habit formation (Miller et al., 2019) where a tendency to select action *i* in state *j* is also denoted by *H*_*i,j*_. Additionally, we introduce another prior given in Equation 8.2, which ensures that only one action has intensity significantly deviating from 0. Furthermore, to link the framework with classical reinforcement learning, we enforce a third condition ensuring that action intensity remains between 0 and 1 (Equation 8.3). These additional priors will often result in one entry of 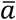 converging to 1, while all other entries decaying towards 0 due to competition. Since in our simulations we also use a binary state vector, the reward expected by the goal-directed system will often be equal to *Q*_*i,j*_ as in the classical reinforcement learning (see paragraph above).

Methods section derives equations describing inference and learning for the above probabilistic model, which for binary state and action vectors take a very similar form to those in standard models of reinforcement learning and a recent model of habit formation (Miller et al., 2019). In particular, at the start of the planning process the dynamics of action intensity is described by Equation 8.4, where the action intensity changes proportionally to a sum of inputs form the goal-directed and habit systems weighted by their confidence, analogously as in a previous model (Miller et al., 2019). The action with a larger initial input is likely to win the competition, so the action for which the right hand side of Equation 8.4 is highest is most likely to be selected. Furthermore, after selection of action *i* in state *j*, the parameters are updated according to Equations 8.5. Namely, the parameter *Q*_*i,j*_ describing expected reward for action *i* in state *j* is modified proportionally to a reward prediction error, as in classical reinforcement learning. Additionally, for every action and current state *j* the parameter describing a tendency to take this action is modified proportionally to a prediction error equal to a difference between the intensity of this action and the intensity expected by the habit system, as in a model of habit formation (Miller et al., 2019).

The similarity of a model developed in the DopAct framework to classical reinforcement learning, which has been designed to maximize resources, highlights that the model also tends to maximize resources, when animal’s reserves are sufficiently low. But the framework is additionally adaptive to the levels of reserves: If the reserves were at the desired level, then *R* = 0 during action planning, so according to Equation 8.4, the goal-directed system would not suggest any action.

Weighting the contribution of the goal-directed system by *R* in Equation 8.4 also has another function: it can bring the contributions of the two systems to the same range, irrespectively of the magnitudes of reward used in the task. Note that in the orange term of Equation 8.4, both *R* and ***Q*** have units of reward magnitude, while Σ_*g*_ has units of reward magnitude squared (because it is a variance), hence the whole orange term is unit-less. The blue term in Equation 8.4 corresponding to the habit system is also unit-less. Therefore, if a task has stochastic rewards, and their magnitude is scaled up, the value of the orange term will not change to a different range, which allows the habit system to contribute to the action selection process irrespective of the scale of rewards.

In the case of deterministic rewards or behaviour, the variance parameters may approach 0 due to learning over trials, which would result in both terms in Equation 8.4 diverging to infinity. To prevent this from happening, a constraint (or a “hyperprior”) on the minimum value of the variance parameters needs to be introduced (in all simulations, if Σ_*g*_ or Σ_*h*_ decreased below 0.2, it was set to 0.2).

The Methods section describes how the inference and learning can be implemented in a generalized version of the network described above. In this network, striatum, output nuclei and thalamus included neural populations selective for the two alternative actions, as in standard models of action selection in the basal ganglia (Bogacz & Gurney, 2007; Frank, Samanta, Moustafa, & Sherman, 2007; Gurney, Prescott, & Redgrave, 2001). The prediction error in the habit system (blue Equation 8.5) is a vector, so computing it explicitly would also require multiple populations of dopaminergic neurons in the habit system selective for available actions, but different dopaminergic neurons in the real brain may not be selective for different actions (da Silva et al., 2018). Nevertheless, this vector has a characteristic structure with large redundancy, so it was sufficient for the simulated model to include only a single population computing the prediction error for the chosen action.

To illustrate predictions made by the model, we simulated it in a probabilistic reversal task. On each trial, the model was “presented” with one of two “stimuli”, i.e. one randomly chosen entry of vector 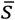 was set to 1, while the other entry was set to 0. On the initial 150 trials, the correct response was to select action 1 for stimulus 1 and action 2 for stimulus 2, while on the subsequent trials, the correct responses were reversed. The mean reward was equal to 1 for a correct response and 0 for an error. In each case, a Gaussian noise with standard deviation *σ*_*r*_ = 0.5 was added to the reward.

Figure 9A shows changes in action intensity and inputs from goal-directed and habit systems as a function of time on different trials within a simulation. On an early trial (left display) the changes in action intensity were primarily driven by the goal-directed system. The intensity of the correct action converged to 1, while it stayed at 0 for the incorrect one. After substantial training (middle display), the changes in action intensity were primarily driven by the habit system. Following a reversal (right display) one can observe a competition between the two systems: Although the goal-directed system had already learned the new contingency (solid orange curve), the habit system still provided larger input to the incorrect action node (dashed blue curve). Since the habit system was faster, the incorrect action had higher intensity initially, and only with time, the correct action node received input from the goal-directed system, and inhibited the incorrect one.

**Figure 9.**
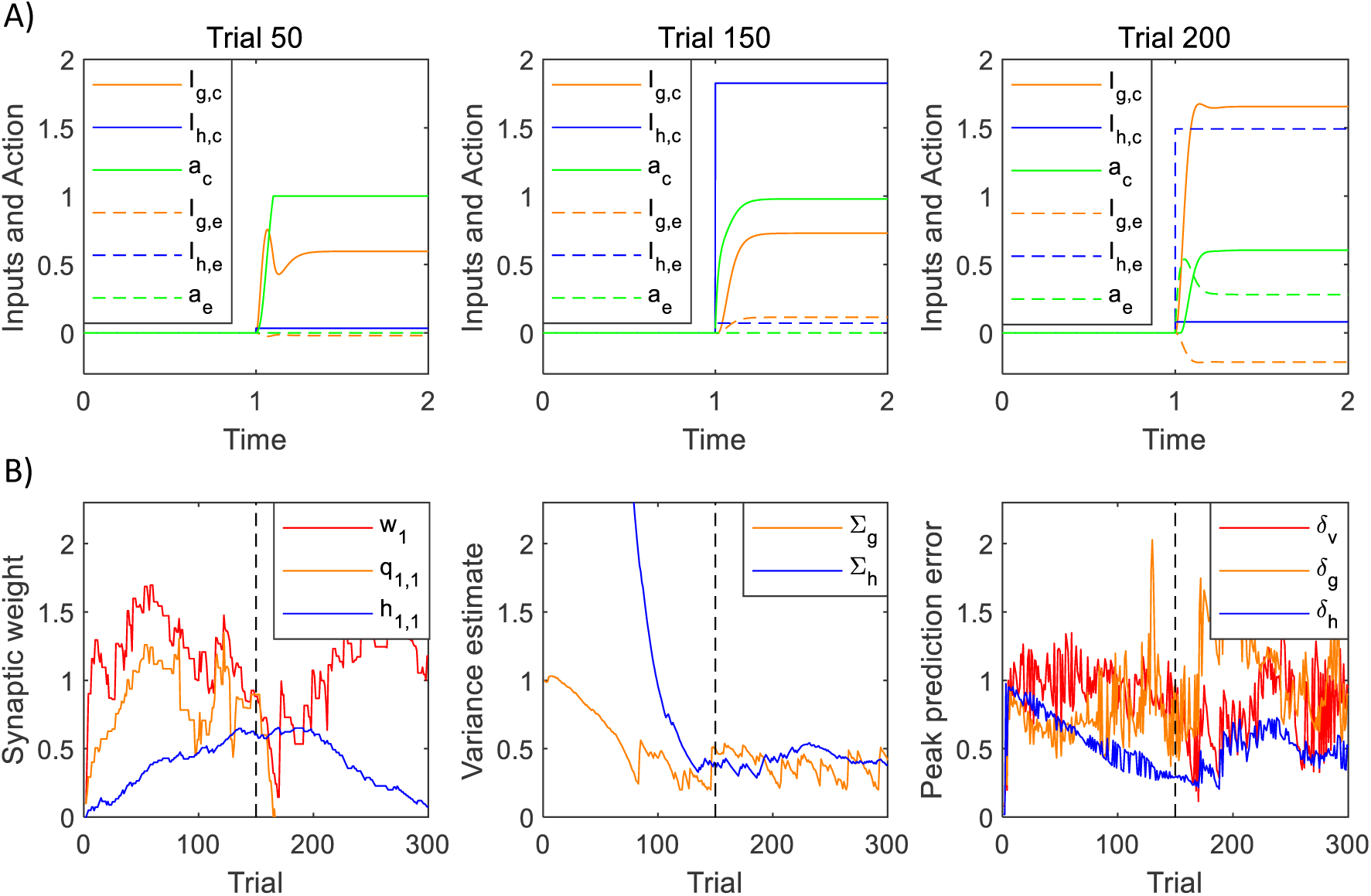
Simulation of the model of choice between two actions. A) Changes in action intensity and inputs from the goal-directed and habit systems, defined below Equation 8.4. Solid lines correspond to a correct action and dashed lines to an error. Thus selecting action 1 for stimulus 1 (or action 2 for stimulus 2) correspond to solid lines in left and middle display (before reversal) and to dashed lines in the right display (after reversal). B) Changes in model parameters and prediction errors across trials. Dashed black lines indicate a reversal trial.

Figure 9B shows how parameters and prediction errors in the model changed over trials. Left display illustrates changes in sample cortico-striatal weights in the three systems. The valuation system rapidly learned the value available after the first stimulus, but after reversal this estimate decreased, as the model persevered in choosing the incorrect option. Once the model discovered the new rule, the estimated value of the stimulus increased. The goal-directed system learned that selecting the first action after the first stimulus gave higher rewards before reversal, but not after. The changes in the parameters of the habit system followed those in the goal-directed system. The middle display shows that the variance estimated by the habit system initially decreased, but then increased several trials after the reversal, when the goal-directed system discovered the new contingency, and thus selected actions differed from the habitual ones. The right display shows an analogous pattern in dopaminergic activity, where the neurons in the habit system signalled higher prediction errors following a reversal. The prediction errors in the goal-directed system increased in a period after reversal, when the agent started to explore the option that had been unrewarded before reversal but now gave higher rewards. These simulations illustrate several experimental predictions of the model to which we will come back in Discussion.

## Discussion

In this paper, we proposed how an action can be identified through Bayesian inference, where the habit system provides a prior and the goal-directed system represents reward likelihood. Within the DopAct framework, the goal-directed and habit systems may not be viewed as fundamentally different systems, but rather as analogous segments of neural machinery performing inference in a hierarchical probabilistic model (Figure 1B), which correspond to different levels of hierarchy.

In this section, we discuss the relationship of the framework to other theories and experimental data, and suggest experimental predictions and directions for future theoretic work.

### Relationship to other theories

The DopAct framework combines elements from four theories: reinforcement learning, active inference, habit formation, and planning as inference. For each of the theories we summarize similarities, and highlight the ways in which the DopAct framework extends them.

As in classical reinforcement learning (Houk et al., 1995; Montague et al., 1996), in the DopAct framework the dopaminergic neurons in the valuation and goal-directed systems encode reward prediction errors. Furthermore, similarly to reinforcement learning models, the parameters describing expected reward are encoded in cortico-striatal weights. The learning rules for these parameters aim at minimizing prediction errors over time, and involve tri-factor Hebbian plasticity often used in models of reinforcement learning (Frémaux & Gerstner, 2016; Kuśmierz et al., 2017; Roelfsema & Holtmaat, 2018). However, the key conceptual difference of the DopAct framework is that it assumes that the goal of animals’ behaviour is to achieve a desired level of reserves, rather than always maximize acquiring resources. It has been proposed that when a physiological state is considered, the reward an animal aims to maximize can be defined as a reduction of distance between the current and desired levels of reserves (Juechems & Summerfield, 2019; Keramati & Gutkin, 2014). Under this definition, a resource is equal to such subjective reward only if consuming it would not bring the animal beyond its optimal reserve level. When an animal is close to the desired level, acquiring a resource may even move the animal further from the desired level, resulting in a negative subjective reward. As the standard reinforcement learning algorithms do not consider physiological state, they do not always maximize the subjective reward defined in this way. Nevertheless, we highlighted (Figure 8), that when the level of reserves is low, the framework can produce a similar behaviour to standard reinforcement learning models (before an action becomes habitual), but importantly, the framework offers flexibility to stop acquiring resources, when the reserves reach the optimum level.

The DopAct framework relies on key high-level concepts from the active inference theory (Buckley et al., 2017; Friston, 2010): the animals aim to reach the desired level of reserves, prediction errors can be minimized by both learning and action planning, and both of these processes can be derived from minimization of free-energy. In the DopAct framework, the neurons encoding prediction errors affect both the plasticity and the activity of its target neurons, analogously as in previous predictive coding architectures derived through free-energy minimization (Friston, 2005). In addition to being a conceptual model, active inference also describes a mechanistic model computing the optimal action (Friston et al., 2013), but the details of the neural implementation of active inference in that model are very different than in DopAct framework. Nevertheless, the function of dopamine is related to that proposed in a past active inference model: in that model it encodes precision, i.e. an inverse variance of choice policy (Friston et al., 2013), while in the DopAct framework it encodes a precision weighted prediction error in the goal-directed system.

We have demonstrated that for a certain set of assumptions, the DopAct framework describes choice and learning processes in a very similar way to a recent model of habit formation (Miller et al., 2019). Both in that model and the DopAct framework, the parameters of the habit system are updated on the basis of prediction errors that do not depend on reward, but rather encode the difference between the chosen and habitual actions. The simulations of several paradigms in this paper parallel those in a previous study (Miller et al., 2019). The key new contribution of this paper is to offer a normative rationale for how such model can arise from Bayesian inference in which the habit system provides a prior. Furthermore, we proposed how learning in this model can be implemented in the basal ganglia circuit including multiple populations of dopaminergic neurons encoding different prediction errors.

Similarly as in the model describing goal-directed decision making as probabilistic inference (Solway & Botvinick, 2012), the actions selected in the DopAct framework maximize a posterior probability of action given the reward. Our simulations of behaviour in devaluation experiments were inspired by the simulations in that study. The new contribution of this paper is making explicit the rationale for why such probabilistic inference is the right thing for the brain to do: The resource that should be acquired in a given state depends on the level of reserves, so the inferred action should depend on what portion of reward available is necessary to restore the reserves. We also propose a detailed implementation of the probabilistic inference in the basal ganglia circuit.

It is useful to discuss the relationship of the DopAct framework to several other theories. It has been suggested that action planning involves a competition between model-based and model-free systems, which are located in prefrontal cortex and striatum respectively (Daw, Niv, & Dayan, 2005). Here we propose that even within the striatum there are circuits maintaining a simple model of how rewards depend on actions. The paper describing a theory of habit formation (Miller et al., 2019) contains an excellent and extensive discussion of its relationship to model-based and model-free theory. Due to the similarity of computations in DopAct framework and in the model of habit formation (Miller et al., 2019), this discussion also applies to the DopAct framework, and we refer interested readers to it. The tonic level of dopamine has been proposed to determine the vigour of movements (Niv, Daw, Joel, & Dayan, 2007). In our model selecting action intensity, the dopaminergic signals in the valuation and goal-directed systems indeed influence the resulting intensity of movement, but in the DopAct framework, it is the phasic rather than tonic dopamine that determines the vigour, in agreement with recent data (da Silva et al., 2018). It has been also proposed that dopamine encodes incentive salience of the available rewards (Berridge & Robinson, 1998; McClure, Daw, & Montague, 2003). Such encoding of incentive salience is present in the DopAct framework, where the prediction error in the goal-directed system depends on whether the available resource is desired by an animal.

### Relationship to experimental data

Relating DopAct framework to experimental data is not fully straightforward for two reasons. First, the tasks simulated in this paper are much simpler than the tasks studied in experiments, in which stimuli and actions have multiple dimensions, and animals often need to make predictions about temporal effects of their behaviour. To fully account for a wide repertoire of neural responses observed in these tasks, more complex models would need to be developed and trained within the DopAct framework. Nevertheless, in this section we will extrapolate and relate the qualitative patterns in presented simulations with available data. Second, to relate the simulations with data, we will need to assume a particular mapping of different systems on anatomically defined brain regions. Thus we will assume that dopaminergic neurons in valuation, goal-directed, and habit systems can be mapped on a spectrum of dopaminergic neurons ranging from ventral tegmental area (VTA) including the valuation system to substantia nigra pars compacta (SNc) including the habit system. The mapping of the dopaminergic neurons from the goal-directed system is less clear, so let us assume that these neurons may be present in both areas. Furthermore, we will assume that the striatal neurons in valuation, goal-directed, and habit systems can be approximately mapped on ventral, dorsomedial, and dorsolateral striatum. However, the neurons corresponding to different systems may not be perfectly separated in space. Keeping these limitations in mind, let us consider the relationship of the DopAct framework to data on functional anatomy, physiology and behaviour.

In the DopAct framework the role of dopamine during action planning is specific to preparing goal-directed but not habitual movements (Figure 6B right). Thus the framework is consistent with an observation that blocking dopaminergic transmission slows responses to reward-predicting cues early in training, but not after extensive training, when the responses presumably became habitual (Choi et al., 2005). Analogously, the DopAct framework is consistent with an impairment in Parkinson’s disease for goal-directed but not habitual choices (de Wit, Barker, Dickinson, & Cools, 2011) or voluntary but not cue driven movements (Johnson et al., 2016). The difficulty in movement initiation in Parkinson’s disease seems to depend on whether the action is voluntary or in response to a stimulus, so even highly practiced movements like walking may be difficult if performed voluntarily, but easier in response to auditory or visual cues (Rochester et al., 2005). Such movements performed to cues are likely to engage the habit system, because responding to stimuli is a hallmark of habitual behaviour (Dickinson & Balleine, 2002).

Dopaminergic modulation of plasticity is required for forming habits in the DopAct framework. Thus the framework is consistent with the observation that lesions to SNc (which we assumed to contain neurons encoding *δ*_*h*_) prevents habit formation (Faure, Haberland, Condé, & El Massioui, 2005). In that study, the lesioned animals could learn to perform an action leading to reward, but after extensive training, they performed action after reward devaluation less frequently than the control animals (implying an impairment in habit formation).

The mapping of the goal-directed and habit systems on dorsomedial and dorsolateral striatum is consistent with the observation that deactivation of posterior dorsomedial striatum impairs learning which action leads to larger rewards (Yin, Ostlund, Knowlton, & Balleine, 2005), while lesion of dorsolateral striatum prevents habit formation (Yin, Knowlton, & Balleine, 2004). The mapping of systems from DopAct framework on subdivisions of the striatum is also consistent with the pattern of neural activity in the striatum, which shifts from encoding reward expectation to movement as one progresses from ventral to dorsolateral striatum (Burton et al., 2015), and with increased activity in dorsolateral striatum during habitual movements (Tricomi, Balleine, & O’Doherty, 2009).

Let us now discuss the relationship of DopAct framework to responses of dopaminergic neurons. These responses have been recorded in different behavioural paradigms, which we discuss in turn. In classical conditioning, dopaminergic neurons in VTA have been shown to encode reward prediction error (Eshel et al., 2016; Schultz et al., 1997; Tobler et al., 2005). Since in the DopAct framework the valuation system is similar to the standard temporal difference learning model, it inherits the ability to account for the dopaminergic responses to unexpected rewards previously explained with that model. It is intriguing to ask if some of the observed dopaminergic responses to a conditioned stimulus reflect the prediction error in the goal-directed system. The motivation for this question is that even in classical conditioning tasks, the animal needs to perform some actions to consume the reward, e.g. swallow it, and it has been reported that animals start to lick after the conditioned stimulus (Tobler et al., 2005). To answer the above question, one would need to analyse how the responses to the conditioned stimulus change across trials. According to the DopAct framework, dopaminergic responses in the goal-directed system should diminish as the action becomes habitual (Figure 6C right). Thus if there existed dopaminergic neurons that encoded reward prediction error in early stages of classical conditioning, but diminished in later stages, then the DopAct framework would suggest they are a part of the goal-directed system.

In operand conditioning tasks, dopaminergic responses to movements performed in order obtain a reward were observed in both VTA and SNc (Engelhard et al., 2019; Schultz, 1986). Analogous dopaminergic responses in all systems within DopAct framework were produced in simulations (Figure 6B left). At the time of reward delivery, the DopAct framework predicts a response in valuation and goal-directed systems, but not in the habit system. Accordingly, a much larger fraction of neurons has been reported to respond to reward in VTA (Engelhard et al., 2019) than in SNc (Schultz, 1986). The DopAct framework also predicts that the responses to movements should be modulated by reward magnitude in the valuation and goal-directed systems, but not in the habit system. This prediction can be compared with data from a task in which animals could press one of two levers that differed in magnitude of resulting rewards (Jin & Costa, 2010). As mentioned above, the framework predicts that the dopaminergic neurons in the valuation and goal-directed systems would respond differently depending on which lever was pressed, while the dopaminergic neurons in the habit system would have response dependent just on action intensity but not reward magnitude. Indeed, a diversity of dopaminergic neurons have been observed in SNc, and the neurons differed in whether their movement related response depended on reward available (Figure 4j in the paper by Jin and Costa (2010)).

The dopaminergic responses have also been observed in a task in which mice could make spontaneous movements and reward was delivered at random times (Howe & Dombeck, 2016). It has been observed that a fraction of dopaminergic neurons had increased responses to rewards, while a group of neurons responded to movements. Moreover, the reward responding neurons were located in VTA while most movement responding neurons in SNc (Howe & Dombeck, 2016). In that study the rewards were delivered to animals irrespectively of movements, so the movements they generated were most likely not driven by processes aiming at achieving reward (simulated in this paper), but rather by other inputs (modelled by noise in our simulations). To relate this task to the DopAct framework, let us consider the prediction errors likely to occur at the times of reward and movement. At the time of reward the animal was not able to predict it, so *δ*_*ν*_ > 0, *δ*_*g*_ > 0, but it was not necessarily making any movements *δ*_*h*_ = 0, while at the time of a movement the animal might have not expected reward *δ*_*ν*_ = *δ*_*g*_ = 0, but might have made non-habitual movements *δ*_*h*_ > 0. Hence the framework predicts separate groups of dopaminergic neurons to produce responses at times of reward and movements, as experimentally observed (Howe & Dombeck, 2016). Furthermore, the peak of the movement related response of SNc neurons was observed to occur after the movement onset (Howe & Dombeck, 2016), which suggests that most of this dopaminergic activity was a response to a movement rather than a response initiating a movement. This timing is consistent with the role of dopaminergic neurons in the habit system, which compute a movement prediction error, rather than initiate movements.

In tasks involving both operand conditioning and spontaneous movements, it has been observed that a fraction of dopaminergic neurons in SNc pause during movements (Dodson et al., 2016; Schultz et al., 1983). It is intriguing to ask if these decreased responses could correspond to brief decreases in activity of dopaminergic neurons in the habit system present during simulations of habitual movements (Figure 6B middle). This may not be a case, because the decreases in simulation are very brief, while they could last ~200ms in SNc neurons (Dodson et al., 2016). Furthermore, a recent study suggests that the decreases of activity of dopaminergic neurons specifically occur during brief movements (“jerks”) that do not develop into longer movements (Howe et al., 2019), and the difference between movement duration was not modelled in the presented simulations. Nevertheless, further investigation of pauses in dopaminergic activity in relationship to DopAct framework may be an interesting direction of future work.

The DopAct framework is consistent with behavioural data connected with habit formation. Analogously to a previous model of habit formation (Miller et al., 2019), it accounts for the effects of training duration and protocol on the perseveration in omission and devaluation paradigms (Figure 7). Additionally, the framework describes the dynamics of competition between the systems during action planning. In a recent study, human participants were extensively trained to make particular responses to given stimuli (Hardwick, Forrence, Krakauer, & Haith, 2018). After a reversal, they tended to produce incorrect habitual actions when required to respond rapidly, but were able to produce the correct actions given sufficient time. Analogous behaviour of the model is shown in the right display of Figure 9A, where the faster habit system initially prepares an incorrect action, but later the slower goal-directed system increases the intensity of the correct action.

Furthermore, the DopAct framework accounts for the observation that animals are less likely to persevere after reversal if larger rewards are used (Theios & Blosser, 1965). An important detail of this experiment was that the rewards were deterministic and of constant magnitude within conditions. Such effect of deterministic reward magnitude on reversals is consistent with the model, because the magnitude of the available reward scales the contribution of the goal-directed system (Equation 8.4), but with deterministic rewards, term Σ_*g*_ that normalizes the goal-directed contribution is little affected by reward magnitude (it may stay at its lower boundary determined by a hyperprior). Thus with higher deterministic rewards, the goal-directed system is more likely to overcome the tendency of the habit system to persevere following a reversal.

### Experimental predictions

The analysis in the previous paragraph suggests that the effect of reward magnitude on the tendency to persevere after reversal (Theios & Blosser, 1965) is a result of using deterministic rewards. Thus the DopAct framework predicts that if this study were repeated with stochastic rewards (typical for a natural environment), the contribution of the goal-directed system (Equation 8.4) would be correctly normalized by the variance, and the effect would disappear.

The DopAct framework predicts distinct patterns of activity for different populations of dopaminergic neurons. Dopaminergic neurons in the habit system should respond to movements more, when they are not habitual, e.g. at an initial phase of task acquisition or after a reversal (Figure 9C). When the movements become highly habitual, these neurons should tend to more often produce brief decreases in response (Figure 6B). Furthermore, when the choices become mostly driven by the habit system (as was the case in the simulation of Figure 6, where the variance of habit system decreased much below the variance of the goal-directed system - middle display in panel C), then dopaminergic neurons in the goal-directed system should no longer signal reward prediction error (Figure 6C, right). By contrast, the dopaminergic neurons in the valuation system should signal reward prediction error irrespectively of the stage of task acquisition (Figure 6C, right).

A central feature of the DopAct framework is that the expectation of the reward in the goal-directed system arises from forming a motor plan to obtain it. Thus the framework predicts that the dopaminergic responses in the goal-directed system to stimuli predicting a reward should last longer if planning actions to obtain the reward takes more time. One way to test this prediction would be to optogenetically block striatal neurons expressing D1 receptors in the goal-directed system for a fixed period after the onset of a stimulus, so the action plan cannot be formed. The framework predicts that such manipulation should prolong the response of dopaminergic neurons in that system.

In the DopAct framework dopaminergic neurons increase the gain of striatal neurons during action planning, only in the goal-directed but not in the habit system. Therefore, the framework predicts that the dopamine concentration should have a larger effect on the slope of firing-Input curves for the striatal neurons in the goal-directed than the habit system. This prediction may seem surprising, because striatal neurons express dopaminergic receptors throughout the striatum (Huntley, Morrison, Prikhozhan, & Sealfon, 1992). Nevertheless, it is consistent with reduced effects of dopamine blockade on initiation of habitual movement (Choi et al., 2005) that are known to rely on dorsolateral striatum (Yin et al., 2004). Accordingly, the DopAct framework predicts that the dopaminergic modulation of dorsolateral striatum should primarily affect plasticity rather than excitability of the striatal neurons.

Patterns of prediction errors expected from the DopAct framework could also be investigated with fMRI. Models developed within the framework could be fitted to behaviour of human participants performing choice tasks. Such models could then generate patterns of different prediction errors (*δ*_*ν*_, *δ*_*g*_, *δ*_*h*_) expected on individual trials. Since prediction errors encoded by dopaminergic neurons are also correlated with striatal BOLD signal (O’Doherty et al., 2004), one could investigate if different prediction errors in the DopAct framework are correlated with BOLD signal in different striatal regions.

## Directions for future work

This paper described a general framework for understanding the function of dopaminergic neurons in the basal ganglia, and presented simple models capturing only a subset of experimental data. To describe responses observed in more realistically complex tasks, models could be developed following a similar procedure as in this paper. Namely, a probabilistic model could be formulated for a task, and a network minimizing the corresponding free-energy derived, simulated and compared with experimental data. This section highlights key experimental observations the models described in this paper are unable to capture, and suggests directions for developing models consistent with them.

The presented models do not mechanistically explain the dependence of dopamine release in ventral striatum on motivational state such as hunger or thirst (Papageorgiou, Baudonnat, Cucca, & Walton, 2016). To reproduce these activity patterns, it will be important to extend the framework to describe the computations in the valuation system.

The models do not describe how the striatal neurons distinguish whether dopaminergic prediction error signal should affect their plasticity or excitability, and for simplicity, in the presented simulations we separated in time the planning and learning processes. However, as illustrated in Figure 2, the same dopaminergic signal may need to trigger plasticity in one group of striatal neurons (selective for a past action), and changes in excitability in another group (selective for a future action). It will be important to further understand the mechanisms which can be employed by striatal neurons to appropriately react to dopamine signals (Berke, 2018).

The models presented in this paper described only a part of the basal ganglia circuit, and it will be important to include also other elements of the circuit. In particular, this paper focussed on a subset of striatal neurons expressing D1 receptors, which project directly to the output nuclei and facilitate movements, but another population expressing D2 receptors projects via an indirect pathway and inhibits movements (Kravitz et al., 2010). Several theories have been proposed for the function of these two classes of neurons. For example, it has been suggested that D1 neurons encode the payoffs of actions, while the D2 neurons encode their costs (Collins & Frank, 2014; Möller & Bogacz, 2019). Consequently, it would be interesting to investigate if the framework can be extended so that the mean expected reward represented by the goal-directed system explicitly includes costs such as effort (e.g. mean reward is assumed to follow non-monotonic functions of action intensity like that in Figure 6A). It could be then investigated if the inference in such modified probabilistic model could be mapped on a network including the striatal D2 neurons. Furthermore, it has been shown how the D1 and D2 neurons can jointly encode the variability of rewards associated with selecting particular actions in a given state (Mikhael & Bogacz, 2016). It would be interesting to investigate if that model can be used to extend the framework so that the variance of reward distribution in the goal-directed system is learned for individual actions.

The basal ganglia circuit also includes a hyperdirect pathway, which contains the subthalamic nucleus. It has been proposed that a function of the subthalamic nucleus is to inhibit non-selected actions (Gurney et al., 2001), and the hyperdirect pathway may support the competition between the actions present in the framework. It will be important to investigate how the definition of the additional priors ensuring that only a single action is selected (Equation 8.2) can be extended to a choice between multiple actions to effectively supress multiple non-selected actions, and how enforcing of such a prior may be achieved by the basal ganglia circuit. The subthalamic nucleus has also been proposed to be involved in determining when the planning process should finish and action should be initiated (Frank et al., 2007). For simplicity, we have simulated the planning process for a fixed interval (Figures 6 and 9), but it will be important to extend the framework to describe the mechanisms initiating an action.

The presented models cannot reproduce the gradual ramping of activity of dopaminergic neurons, observed as animals approached rewards (Howe, Tierney, Sandberg, Phillips, & Graybiel, 2013). To be consistent with these data, the valuation system could be extended to incorporate synaptic decay that has been shown to allow standard reinforcement learning models to reproduce the ramping of prediction error (Kato & Morita, 2016). The described models assume that there are only 3 groups of dopaminergic neurons with 3 types of responses. To model further diversity of dopaminergic responses within each system (Dodson et al., 2016; Jin & Costa, 2010) the framework could be extended to assume that each system tries to predict multiple dimensions of reward or movement (cf. Gardner, Schoenbaum, & Gershman, 2018).

The models did not describe a general tendency for making movements when reward becomes available, known as Pavlovian biases (Guitart-Masip et al., 2012). To incorporate such Pavlovian biases in the DopAct framework, it can be assumed that the mean of reward expected by the goal-directed system also depends directly on the action (e.g. one could modify the model selecting action intensity so that the mean expected reward is given by *aqs* + *ba*, where *b* is a parameter describing the strength of Pavlovian biases).

Finally, this paper focusses on the role of dopamine in the basal ganglia. Dopaminergic neurons additionally project to the cortex, where they have been proposed to modulate synaptic plasticity (Roelfsema & Ooyen, 2005). However, dopaminergic neurons also modulate excitability of cortical neurons (Thurley et al., 2008), so it would be interesting to investigate if they can trigger action planning by cortical networks.

## Methods

This section describes details of models developed within the DopAct framework for two tasks: selecting action intensity and choice between two actions. The models were simulated in Matlab, and all codes are available at MRC Brain Network Dynamics Unit Data Sharing Platform (https://data.mrc.ox.ac.uk/data-set/simulations-action-inference).

### Selecting action intensity

We first describe an extended version of the actor that learns uncertainties associated with goal-directed and habit systems, and then provide the details of the dynamics of the simulated model.

Substituting probability densities of likelihood and prior distributions (Equations 3.2-3.3) into Equation 4.1, we obtain an expression for the objective function *F* in Equation 10.1 in Figure 10A, which has an analogous form as before but now includes the variance parameters. To find action intensity, we change it according to the gradient of *F*, as described in Equation 10.2. Analogously as in the simple case described in the Results, the action intensity is driven by both goal-directed and habit systems, but now their contributions are normalised by the variance parameters. For the habit system this normalization is stated explicitly in Equation 10.2, while for the goal-directed system it comes from a modified definition of prediction error in orange Equation 10.3. Since variance of the habit system already normalizes its contribution in Equation 10.2, we choose not to include it in the definition of habit prediction error.

**Figure 10.**
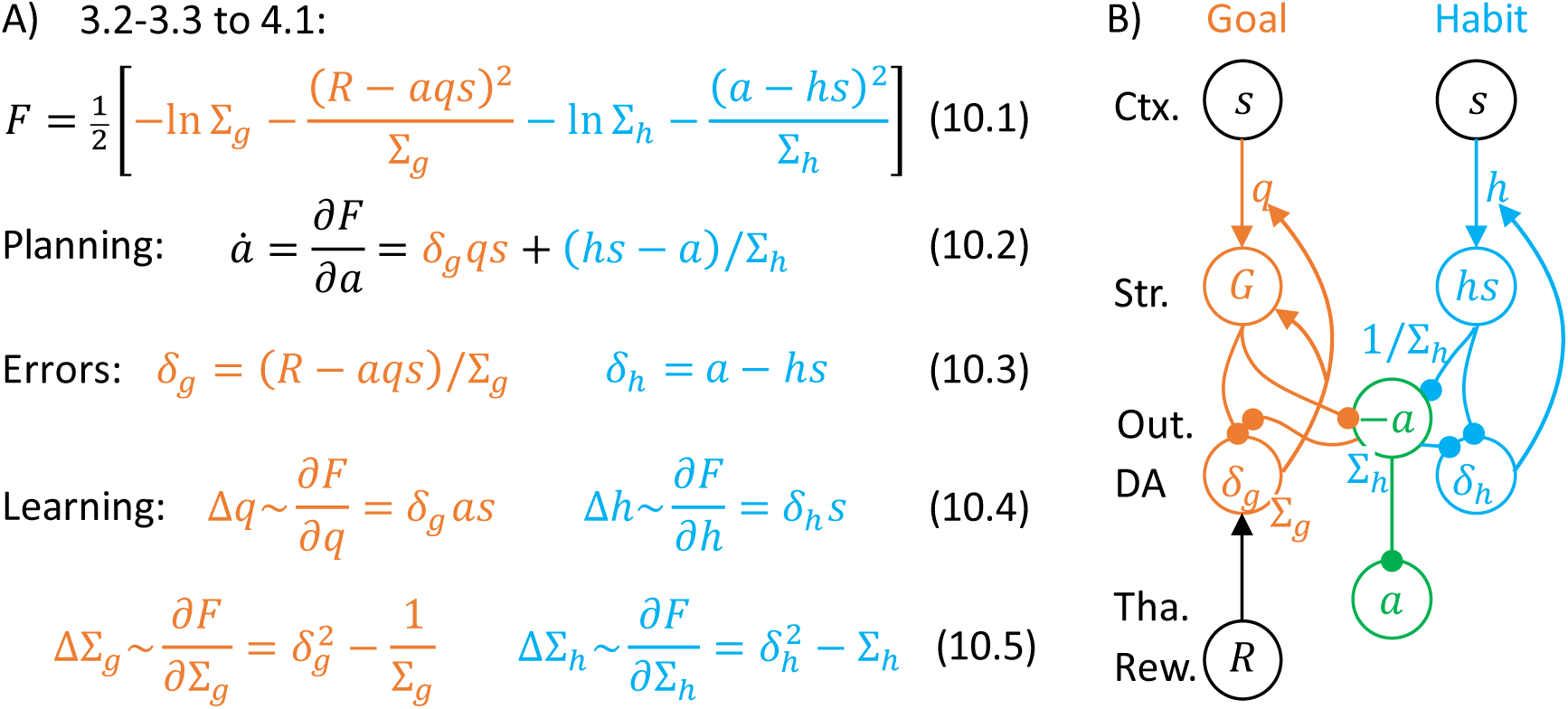
Description of a model selecting action intensity. A) Details of the algorithm employed in the model. B) Mapping of the algorithm on network architecture. Notation as in Figure 5B.

When a reward is obtained, the parameters are modified proportionally to the derivatives of *F* over the parameters. The update rules for parameters describing mean of the distribution in Equation 10.4 remain the same as in Equation 5.4 (ignoring constants that can be incorporated into a learning rate). The update rule for the parameter describing variance in the goal-directed system can be obtained by computing derivative of *F*, giving the orange Equation 10.5. Following an analogous procedure for the variance in the habit system, we would obtain a derivative equal to 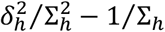 but to simplify this expression, we scale it by 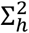, resulting in the blue Equation 10.5. As 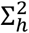 is a positive number, such scaling does not change the value to which Σ_*h*_ converges.

There are several ways of including the variance parameters in the network, and as a proof of principle we show here one of them (Figure 10B). This network is very similar to that shown in Figure 5B, but now the projection to output nuclei from the habit system is weighted by its precision 1 /Σ_*h*_ (to reflect the weighting factor in Equation 10.2), and also the rate of decay (or relaxation to baseline) in the output nuclei could be proportional to Σ_*h*_. One way to ensure that prediction error in goal-directed system is scaled by Σ_*g*_ is to encode Σ_*g*_ in the rate of decay or leak of these prediction error neurons (Bogacz, 2017; Friston, 2005). To see how the scaling of *δ*_*g*_ by Σ_*g*_ in orange Equation 10.3 arises, the dynamics of these prediction error neurons can be written as a differential equation, which is included as orange Equation 11.2 in Figure 11. This equation, includes at the end a decay term with a decay rate Σ_*g*_. To find a value to which orange Equation 11.2 converges, we note that in equilibrium 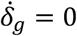, so by setting left hand of the equation to 0 and solving for *δ*_*g*_ we obtain the value of *δ*_*g*_ in an equilibrium. This value is equal to that in orange Equation 10.3, with a difference that the total reward *R* is here replaced by the sum of instantaneous reward *r*, and available reward v provided by the valuation system, which are the inputs to these prediction error neurons (Figure 1D).

**Figure 11.**
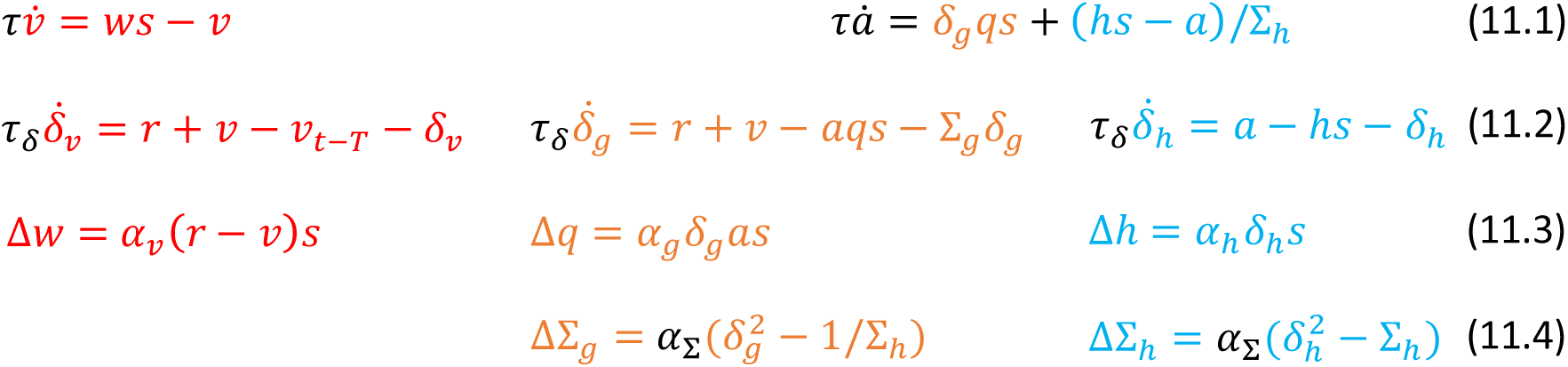
Dynamics of the model choosing action intensity. Red, orange and blue equations describe valuation, goal-directed and habit systems, respectively.

The updates of the variance parameter (Equations 10.5) only depend on the corresponding prediction errors and the variance parameters themselves, so they could be implemented with local plasticity if the neurons encoding variance parameters received corresponding prediction errors.

Figure 11 provides a complete description of the dynamics of the simulated model. The orange and blue equations describe the actor and they parallel those in Figure 10A, but now explicitly include time constants for nodes encoding action intensity (*τ*) and prediction errors (*τ*_*δ*_), as well as learning rates for the means of distributions in the goal-directed (*α*_*g*_) and habit (*α*_*h*_) systems and for variances (*α*_Σ_). Red equations provide a description of the valuation system. According to red Equation 11.1, the estimate of the value of state *s* converges in equilibrium to *ν* = *ws*. To illustrate the dynamic of the prediction error of the valuation system, it is simulated according to red Equation 11.2, which converges to a difference between total reward (*r* + *ν*) and the expectation of that reward made at an earlier time (*ν*_*t-T*_, where *T* is a parameter describing how long ago the prediction was made). This prediction error is similar to that in the standard temporal difference learning (Sutton & Barto, 1998), but for simplicity we do not represent the vector of parameters encoding the reward predicted in different moments in time. Dynamics of such prediction error after conditioned stimulus will be similar to that in a trained temporal difference learning model, where all parameters encoding reward expectation until time it typically occurs converge to the same value (if the “discount factor” is set to 1). This prediction error is used only to illustrate expected dynamics of dopaminergic neurons in the valuation system, but it does not drive plasticity. Following reward delivery, the parameter *w* is modified according to red Equation 11.3, where *ν* is taken as the estimated value at the end of simulation of the planning phase on this trial, and *α*_*v*_ is a learning rate.

We assume that the intensity of action executed by the agent is equal to the inferred action intensity plus motor noise with standard deviation *σ*_*α*_. For simplicity we do not explicitly simulate the dynamics of the model after the delivery of reward *r*, but we fix the value of *a* to intensity of executed action. We compute the prediction errors in the goal-directed and habit system in an equilibrium (Equations 10.3), and update the parameters. In simulations the learning rate of the valuation system was set to *α*_*ν*_ = 0.5 on trials with *δ*_*ν*_ > 0, and to *α*_*ν*_ = 0.1 when *δ*_*ν*_ ≤ 0. Other parameters of the simulation were set to: *τ* = 0.05, *τ*_*δ*_ = 0.02, *α*_*g*_ = *α*_*h*_ = 0.02, *α*_*Σ*_ = 0.01, and *T* = 0.1. The model parameters were initialized to *ν* = *q* = 0.1, *h* = 0, Σ_*g*_ = 1 and Σ_*h*_ = 100.

### Choice between two actions

This section describes a sample neural implementation of inference and learning in a probabilistic model in Equations 8.1-8.3. Substituting probability densities from Equations 8.1 and 8.2 into the objective function of Equation 4.1, we obtain Equation 12.1 in Figure 12A. We chose not to explicitly include the constraint of Equation 8.3 into the objective function, but instead we ensured that the action intensity remained in range between 0 and 1 by setting *a*_*i*_ to one of these values if it exceeded the range during numerical integration.

**Figure 12.**
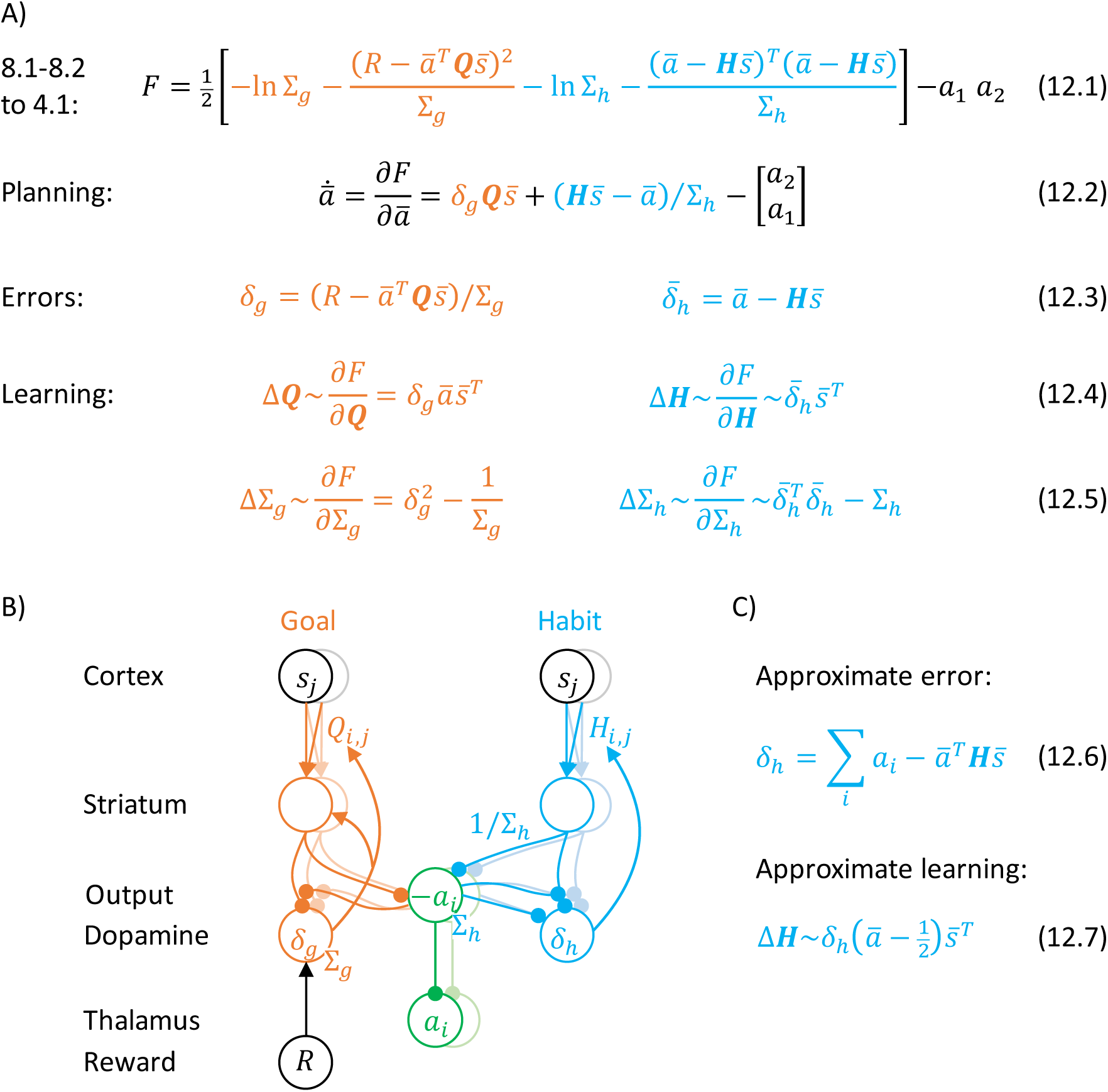
Implementation of the actor in a model of choice between two actions. A) Mathematical description of the algorithm in the model. B) Mapping of the algorithm on network architecture. Notation as in Figure 5B. C) An approximation of learning in the habit system.

To obtain the equations describing action planning or learning, we need to compute derivatives of *F* over vectors or matrices. The rules for computing such derivatives are natural generalizations of the standard rules and they can be found in a tutorial paper (Bogacz, 2017). During planning, the action intensity should change proportionally to a gradient of *F*, which is given in Equation 12.2, where the prediction errors are defined in Equations 12.3. These equations have an analogous form to those in Figure 10A, but are generalized to matrices. The only additional element is the last term in Equation 12.2, which ensures competition between different actions, i.e. *a*_1_ will be decreased proportionally to *a*_2_, and vice versa. During learning, the parameters need also be updated proportionally to the corresponding gradients of *F*, which are given in Equations 12.4 and 12.5. Again, these equations are fully analogous to those in Figure 10A.

It is interesting to note how the equations in Figure 12A simplify in certain relevant limits. First, consider the evolution of action intensity at the start of the trial, when *a*_*i*_ ≈ 0. Substituting orange Equation 12.3 into Equation 12.2 and setting *a*_*i*_ = 0, we obtain Equation 8.4. Second, consider the update rules: if only single elements of vectors 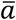 and 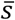 have non-zero values *a*_*i*_ = 1 and *s*_*j*_ = 1, then substituting Equations 12.3 into 12.4 and ignoring constants gives Equations 8.5.

Let us now consider how equations in Figure 12A could be implemented in a neural circuit schematically shown in Figure 12B. There are many ways of achieving these computations, but as a proof of principle, we discuss one candidate mapping. We assume that striatum, output nuclei and thalamus include neural populations selective for the two alternative actions (shown in vivid and pale colours in Figure 12B), and the connections between these nuclei are within the populations selective for a given action, as in previous models (Bogacz & Gurney, 2007; Frank et al., 2007; Gurney et al., 2001). Additionally, we assume that sensory cortex includes neurons selective for different states (shown in black and grey in Figure 12B), and the parameters *Q*_*i,j*_ and *H*_*i,j*_ are encoded in cortico-striatal connections. Then, the orange and blue terms in Equation 12.2 can be computed by the striatal neurons in goal-directed and habit systems in exactly analogous way as in the network inferring action intensity, and these terms can be integrated in the output nuclei and thalamus. The last term in Equation 12.2 corresponds to mutual inhibition between the populations selective for the two actions, and such inhibition could be provided by inhibitory projections that are presents in many different regions of this circuit, e.g. by co-lateral projections of striatal neurons (Preston, Bishop, & Kitai, 1980) or via a subthalamic nucleus, which has been proposed to play role in inhibiting non-selected actions (Bogacz & Gurney, 2007; Frank et al., 2007; Gurney et al., 2001).

The prediction error in the goal-directed system (orange Equation 12.3) could be computed analogously as in the network selecting action intensity as a difference between reward and expectation provided by the striatum. To compute the expected reward 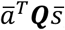, the striatal neurons could calculate 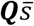 and the elements of this vector could be summed up thanks to projections from striatum to dopaminergic neurons that could be scaled by input encoding 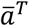 (Figure 12B). During learning, the prediction error in the goal-directed system modulates plasticity of the corresponding cortico-striatal connections according to orange Equation 12.4, which describes a standard tri-factor Hebbian rule (if following movement the striatal neurons encode chosen action, as assumed in Figure 5C). The learning rule for the variance parameter of the goal-directed system (orange Equation 12.5) is exactly the same as in the model in the previous section (cf. orange Equation 10.5).

Learning in the habit system can be approximated with a single dopaminergic population, because the prediction error 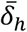 has a characteristic structure with large redundancy. Namely, if the vectors 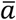 and 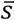 are binary (i.e. only one entry is equal to 1 and other entries to 0), then only one entry in 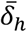 corresponding to the chosen action is positive, while all other entries are negative (because parameters *H*_*i,j*_ stay in a range between 0 and 1 when initialized within this range and updated according to blue Equation 12.4). Hence, we simulated an approximate model with a single prediction error just encoding the prediction error for the chosen action (Equation 12.6). With such a single modulatory signal, the learning rules for striatal neurons in the habit system have to be adjusted so the plasticity has opposite directions for the neurons selective for the chosen and the other actions. Such modified rule is given in Equation 12.7 and corresponds to tri-factor Hebbian learning (if striatal neurons in the habit system have activity proportional to 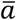 during learning, as we assumed for the goal-directed system). Thanks to this approximation, the prediction error and plasticity in the habit system take a form that is more analogous to that in the goal-directed system. When the prediction error in the habit system is a scalar, the learning rule for the variance parameter (blue Equation 12.5) becomes the same as in the model in the previous section (cf. blue Equation 10.5).

In order to simulate the model, it has been converted to differential equations in analogous way as in the previous section. The simulated model also included a valuation system, which was parameterized by a vector 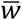 that described the values of the states in the simulation. At the end of the planning phase, Gaussian noise with standard deviation *σ*_*a*_ = 2 was added to all entries of the action vector (to allow exploration), and the action with the highest intensity was “chosen” by the model. Subsequently for the chosen action *i*, the intensity was set to *a*_*i*_ = 1, while for the other action it was set to *a*_*k≠i*_ = 0. All parameters of the simulations had the same value as in the previous section, except for *α*_*g*_ = 0.1 and *α*_*h*_ = 0.05.

## Acknowledgements

This work has been supported by MRC grant MC_UU_12024/5. The author thanks Moritz Moeller and Sashank Pisupati for comments on an earlier version of the manuscript, and Karl Friston, Yonatan Loewenstein, Mark Howe, Friedemann Zenke, Kevin Miller and Peter Dayan for discussion.

